# Assessing the combined performance of supervised learning and spike-in constructs for bias correction in eDNA metabarcoding

**DOI:** 10.1101/2025.03.25.645285

**Authors:** Jason Chang, Rasmus Nielsen

**Affiliations:** Department of Integrative Biology, University of California Berkeley; Berkeley, CA 94720, United States of America; Institute for Computational Medicine, John Hopkins University; Baltimore, MD 21218, United States of America; Department of Biomedical Engineering, John Hopkins University; Baltimore, MD 21218, United States of America; Museum of Vertebrate Zoology, University of California Berkeley; Berkeley, CA 94720, United States of America; Center for Computational Biology, University of California Berkeley; Berkeley, CA 94720, United States of America; Department of Statistics, University of California Berkeley; Berkeley, CA 94720, United States of America; Globe Institute, University of Copenhagen; Copenhagen, 1350, Denmark

## Abstract

Environmental DNA (eDNA) metabarcoding has become increasingly popular as an approach to efficiently document biodiversity within an environment characterized by relative uncertainty. Compared to the traditional stereomicroscopic approaches, eDNA metabarcoding is simpler and less costly. Under ideal circumstances, researchers are able to directly extrapolate the true relative abundance of a particular taxon in the sampled environment by computing the proportion of sequenced reads assigned to the specific taxon. Although several previous studies have been carried out under such assumptions, some researchers have raised the possibility that there may exist both biological and technical biases in eDNA metabarcoding studies, leading to inconsistent estimations of community composition. Using mock community datasets from nine relevant studies in the past, we showed that bias correction in eDNA metabarcoding studies is indeed a predictable task. We also found *reads* and *amp_gc* to be the two most important feature predictors, such that these two features alone are enough to retain most of the model performances. Experiment-specific information were found to be necessary for bias correcting models to perform well. However, we have yet to develop an effective way of converting knowledge regarding spike-in (SP) samples into experiment-specific information that can be learned by existing models. Nonetheless, under the data-specific scenario, AdaBoost showed an optimal 35.62% improvement from the baseline established by the vanilla control model. Additionally, we showed that model performances could be rescued by the availability of experiment-specific data, under which XgBoost exhibited an optimal 81.57% improvement from the baseline. Our work suggests that future metabarcoding studies would benefit from performing supervised learning (SL)-based bias correction prior to downstream analyses. Moreover, if experiment-specific data is available at the time of the study, it is optimal to construct an XgBoost model. Otherwise, it is still recommended to construct an AdaBoost model, which showed marginal improvement from the baseline with no modeling.

**One Sentence Summary:** Supervised learning models, particularly XgBoost and AdaBoost, can effectively correct biases in eDNA metabarcoding studies, with performance improving significantly when experiment-specific data is available.

## Background

DNA metabarcoding is a novel approach used to document biodiversity in environments that lack sufficient information regarding species richness and taxonomic composition given a priori. The simplicity and low cost of DNA metabarcoding has allowed it to become increasingly popular in the field of ecological and palaeoecological research (Nichols et al., 2018). Through external interactions with the surrounding environment, living organisms shed traces of DNA in forms that include but are not limited to skin, scales, hair, and mucus, which essentially accumulate in the surroundings. Such buildup can be extracted as environmental DNA (eDNA) from the environmental source, carrying the genetic information of specific species residing within that ecosystem. The extracted eDNA can subsequently be amplified by polymerase chain reaction (PCR) using universal primers that are complementary to highly conserved marker genes among the target species expected to be found in the sampled environment. The amplicons are then sequenced through next generation sequencing (NGS) and mapped to genome of species in the reference sequence database during sequence alignments.

Theoretically, by computing the proportion of sequenced reads assigned to a taxon, we should be able to extrapolate the true relative abundance of that particular taxon in the sampled environment. However, performing a precise extrapolation is more complicated than it seems. While several previous studies have directly estimated the relative biomass by computing the relative abundance of assigned reads (Evans et al., 2016; Matesanz et al., 2019; Takahara et al., 2012), many researchers have questioned the accuracy of such an approach, citing the factors that are likely to lead to bias and inconsistency when inferring the composition of a microbial community (Brooks et al., 2015; Choo et al., 2015; D’Amore et al., 2016; Deagle et al., 2013; Jones et al., 2015; Mackenzie et al., 2015; Pawluczyk et al., 2015; Piñol et al., 2019).

The potential sources of bias behind the use of relative read abundances as relative biomass estimates can be both biological and technical (Nichols et al., 2018). Biological biases can stem from a variety of factors: the spatial and temporal differences in dispersal of eDNA; the rates at which eDNA is transported and degraded across long distances by environmental media such as air and water; copy number variations in the target loci; and seasonal variations in natural populations (Beentjes et al., 2019; Djurhuus et al., 2020; Krehenwinkel et al., 2017; Krehenwinkel et al., 2018; Loos and Nijland et al., 2020; Sigsgaard et al., 2017; Vasselon et al., 2017). The correction of biological biases primarily depends on the optimization of sampling technique for a given environment. On the other hand, technical biases can be introduced during the extraction, amplification, and sequencing of eDNA templates. eDNA extraction protocols can be suboptimal depending on the types of inhibitory compounds that are present in the environment, which can potentially lead to inconsistency in the genetic material recovered (Hunter et al., 2019; Zielińska et al., 2017). Above all, amplification of eDNA templates is carried out via the PCR, which is in itself a stochastic reaction that critically depends on the thermodynamics and kinetics of polymerase and other reagents involved in the amplification (Pinto and Raskin, 2012). The multiplicity of variable eDNA templates in the pooled experiment further escalates the complexity and inconsistency of the amplification reaction (Polz and Cavanaugh, 1998). Templates of certain amplicon size and GC-content are favored based on the specific PCR profile carried out by the thermocycler (Kumar and Kaur, 2014; Huber et al., 2009). Although universal primers are designed to effectively amplify a wide spectrum of species, the characteristics and positions of internal primer-template mismatches can impose detrimental effects on the primer binding efficiency, introducing more technical biases into the equation (Bru et al., 2008). Lastly, intrinsic NGS error can also emerge from the sequencing process, despite its relative rarity compared to other bias factors (Ma et al., 2019).

A recent solution to the abovementioned bias problem in high-throughput sequencing studies has been the introduction of spike-in control, which constitutes of multiple distinct synthetic oligonucleotides that are not part of the original sample of interest but are added to the sequencing pipeline in order to serve as a calibration standard for downstream analysis of the sequenced reads from a specific experiment. In order to normalize data across multiple experiments using spike-in control, one must first determine the standard spike-in concentration values for normalization. By comparing the normalized spike-in concentrations to the raw read counts of each spike-in, one is able to construct a normalization function that maps from raw read counts to normalized read counts in each experiment (Chen et al., 2016). Formally, let *r_raw_* and *r_norm_* be the raw and normalized read counts of a sample respectively, the normalization function *f* describes a function such that *f(r_raw_)* = *r_norm_*. For clarification, we hereby define a sample of a metabarcoding study as a *(taxon, abundance)* tuple for a specific taxon in one given experiment while a dataset corresponds to a set of mock community experiments that make up the study. In the ideal scenario, the normalization function *f* describes a linear relationship; however, this is not always the case given the complexity of the modern metabarcoding studies. This prompts the need for a more involved model that is able to capture the non-linear relationships between *r_raw_* and *r_norm_*. Better yet, we could potentially construct a model that is robust enough to take in both experimental conditions and *r_raw_* as inputs to directly infer the true abundance *r_true_* of a sample, thereby bypassing the normalization step altogether.

Supervised learning (SL) is a branch of machine learning (ML) problems that aim to construct a concise model of the distribution of object label in terms of predictor features, which is then applied to make predictions regarding future instances (Kotsiantis, 2007). Moreover, classification describes a problem where the label of an object is given in discrete value, indicating which class the object belongs; regression poses a problem where the label of an object is given as a continuous output variable, specifying the quantity associated with the object.

Here, we explore the potential of a correction model that remedies the technical biases introduced in the process of eDNA metabarcoding through the combination of supervised learning and synthetic DNA spike-in constructs. Six SL algorithms, i.e. linear regression, polynomial regression with degree of 2, elastic net regression, AdaBoost, XGBoost, and random forest, were evaluated under different experimental scenarios alongside an identity function as a vanilla control group. XGBoost emerged as the optimally performing model under the scenarios where experiment-specific data is provided; in contrast, AdaBoost was obtained as the optimal model under the scenarios where only global data is given. The two models, XGBoost and AdaBoost, provides two good starting points for correcting bias in future metabarcoding studies depending on the specificity of the data at hand.

## Results

The brief overview of the dataflow and analysis presented in this study are summarized in Fig. 1. For the modeling to be done in the context of simulated environmental samples, we have mined mock community datasets from nine relevant studies (Angel et al., 2018; Bokulich et al., 2015; Callahan et al., 2016; Gohl et al., 2016; Kozich et al., 2013; Leray and Knowlton, 2017; Schirmer et al., 2015; Taylor et al., 2016; Tourlousse et al., 2017). Mock communities consist of defined mixture of microorganisms created *in vitro* to simulate composition of microbial communities found in the wild (Highlander, 2013). For clarification, we hereby refer to the dataset from Tourlousse et al. as the spike-in dataset interchangeably while the remaining studies form the global dataset as Tourlousse et al. were the only group that looked into the performance of synthetic spike-in constructs under metabarcoding studies (2017).

**Figure 1:**
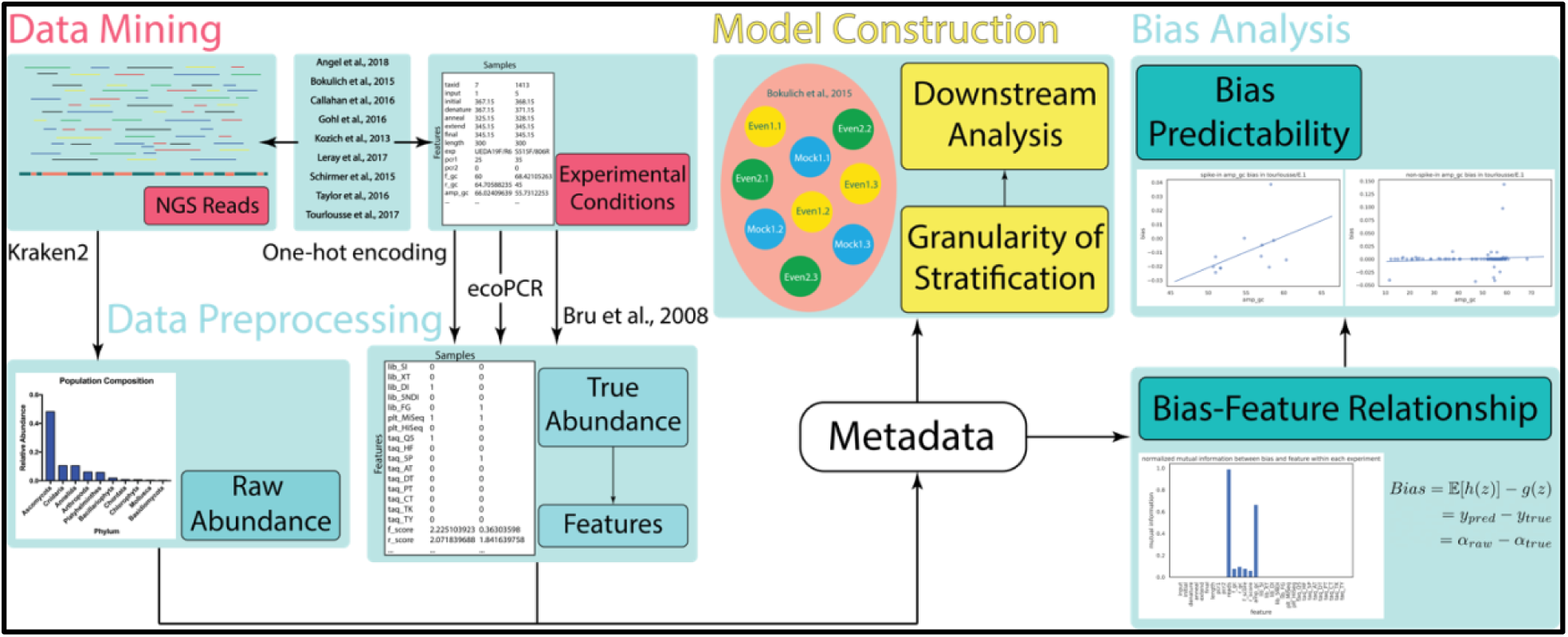
Schematic overview of the dataflow and analysis presented in this study. NGS reads and experimental conditions were mined from the relevant mock community studies. Sequenced reads were taxonomically assigned using the *Kraken2* assignment tool. Experimental conditions were further preprocessed using one-hot encoding to convert categorical variables into quantitative features. Characteristics of primer-template binding and PCR amplification were assessed by *in silico* PCR and primer binding score. The metadata were aggregated for downstream bias analysis and evaluation of model performance.

Information regarding both sequenced reads and experimental conditions were obtained from these datasets. Each dataset represents one prior study, which includes a set of experiments performed in the study. Each unique taxon assigned in an experiment is what we define as a sample in this study. The sequenced reads were taxonomically assigned by *Kraken2* to simulate presumably biased relative abundances from realistic taxonomic assignment workflows. Attributes regarding each experiment were also mined from each literature as feature predictors, such as the temperature at various stages of PCR, sequenced read length, primer GC-contents, total amount of input DNA within each experiment, library preparation method, sequencing platform, and Taq polymerase (Table 1). To analyze the interactions between the primers and the templates closely, we performed *in silico* PCR with the *ecoPCR* pipeline and computed primer binding score for each primer-template matching pair using experimental data provided by Bru et al. in a related study (2008).

**Table 1:** Description of feature predictors included in the extracted dataset. Experimental conditions extracted from each mock community include detailed protocol of the PCR, GC-content of each primer, total amount of input DNA, library preparation method, sequencing platform, and Taq polymerase. Reads here refer to the biased relative abundances from taxonomic assignments outputted by Kraken2. Advanced characteristics regarding the primer-template interactions during the PCR were evaluated by performing *in silico* PCR with the *ecoPCR* pipeline. Scoring of pairwise primer-template binding was extrapolated by a linear model fitted on experimental data published by Bru et al. in a previous study (2008). lib_{X}, plt_{Y}, and taq_{Z} are one-hot encoded features converted from the original categorical values provided in each literature.

| Feature | Description |
| --- | --- |
| input | Total amount of input DNA across all species in the experiment |
| initial | Temperature of initial denaturation in °K |
| denature | Temperature of cycling denaturation in °K |
| anneal | Temperature of cycling annealing process in °K |
| extend | Temperature of cycling elongation in °K |
| final | Temperature of final extension in °K |
| length | Length of the sequenced read in bp |
| pcr1 | Number of cycles in the first PCR reaction |
| pcr2 | Number of cycles in the second PCR reaction (if applicable) |
| reads | Raw relative abundance outputted from Kraken2 within the experiment |
| f_gc | GC-content of the forward primer binding site |
| r_gc | GC-content of the reverse primer binding site |
| f_score | Binding score for the quality of template-forward primer matching |
| r_score | Binding score for the quality of template-reverse primer matching |
| amp_gc | GC-content of the amplicon |
| lib_{X} | Indicator variable of whether the experiment uses library preparation method X, where $X \in \{SI, XT, DI, 5NDI, FG\}$ |
| plt_{Y} | Indicator variable of whether the experiment is sequenced using sequencing platform Y, where $Y \in \{MiSeq, HiSeq\}$ |
| taq_{Z} | Indicator variable of whether the experiment uses Taq polymerase Z, where $Z \in \{Q5, HF, 5P, AT, DT, PT, CT, TK, TY\}$ |

Prior to construction of various regression models, we attempted preliminary analysis of bias to ascertain whether there indeed exist associations between the chosen feature predictors and the bias to be extracted by the ultimate model. By computing the normalized mutual information (NMI) between the bias and each i^th^ feature ( *feature_i_*) vectors within each experiment, we found that observations of *amp_gc* provide the second most information regarding the distribution of bias (Fig. 2). We included *reads* in this analysis as a control group as it is included intrinsically in the derivation of bias, thus it is expected that observations of *reads* would contain the most information regarding quantities of bias. This analysis of NMI prompted the importance of *amp_gc* in the task of bias correction, which we will soon return to when evaluating the importance of each *feature_i_* in performance of various regression models.

**Figure 2:**
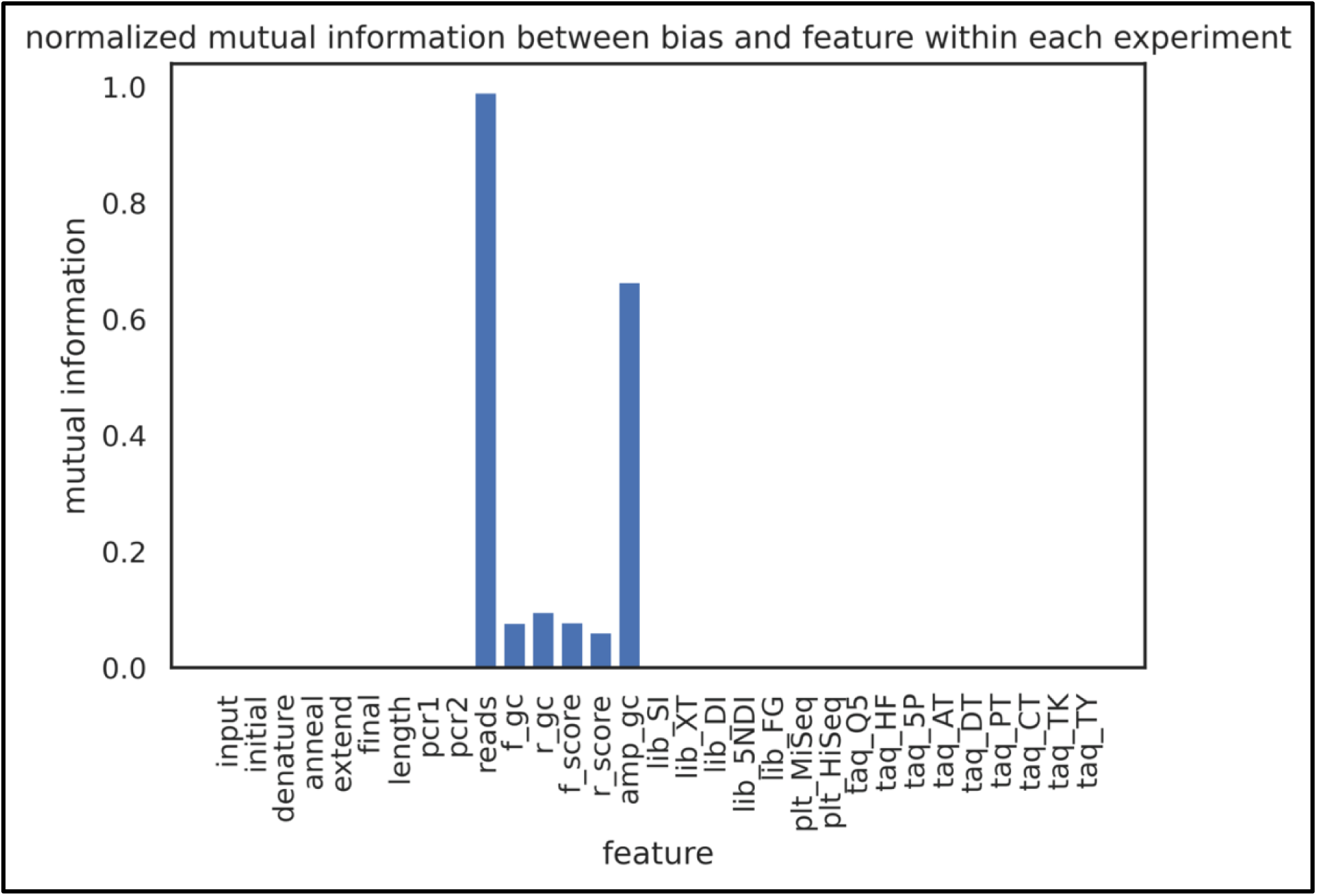
NMI between bias and the i^th^ feature within each experiment. NMI measures the amount of information obtained regarding the distribution of bias from the observation of each *feature_i_*, which ranges in the interval [0, 1] instead of [0, ∞) as in the unnormalized version. *Reads* was included as a control group as it is expected to hold the most information regarding quantity of bias among the features as *reads* is implicitly involved in the derivation of bias.

To examine the potential of synthetic spike-in constructs in bias correction, we first plotted bias against each *feature_i_* for spike-in (SP) and non-spike-in (NSP) samples separately within each experiment of the spike-in dataset (Fig. S1). We then evaluated the correlation between bias and each *feature_i_* by determining the goodness of fit of a linear regression model on the data of each plot, which can be obtained by calculating the mean squared error (MSE) of the linear regression model. Within each experiment, features were ranked based on the MSE for SP and NSP samples individually to form two vectors of ranked features. We computed the Kendall rank correlation coefficient between these two vectors to measure the ordinal association between the SP and NSP ranked features, which would allow us to answer the question of whether strong predictors for SP bias are also strong predictors for NSP bias within the same experiment. We found that this only holds true for a fraction of the experiments in the spike-in dataset when testing for the null hypothesis that the two vectors are ordinally independent of each other, which hinted at the limited strength and specificity of spike-in constructs in bias correction (Fig. 3). The linear regression model is used here for simplicity reason in the preliminary experiment; we understand that perhaps there exists more complicated relationship between bias and each *feature_i_* that can be better observed by increasing model complexity.

**Figure 3:**
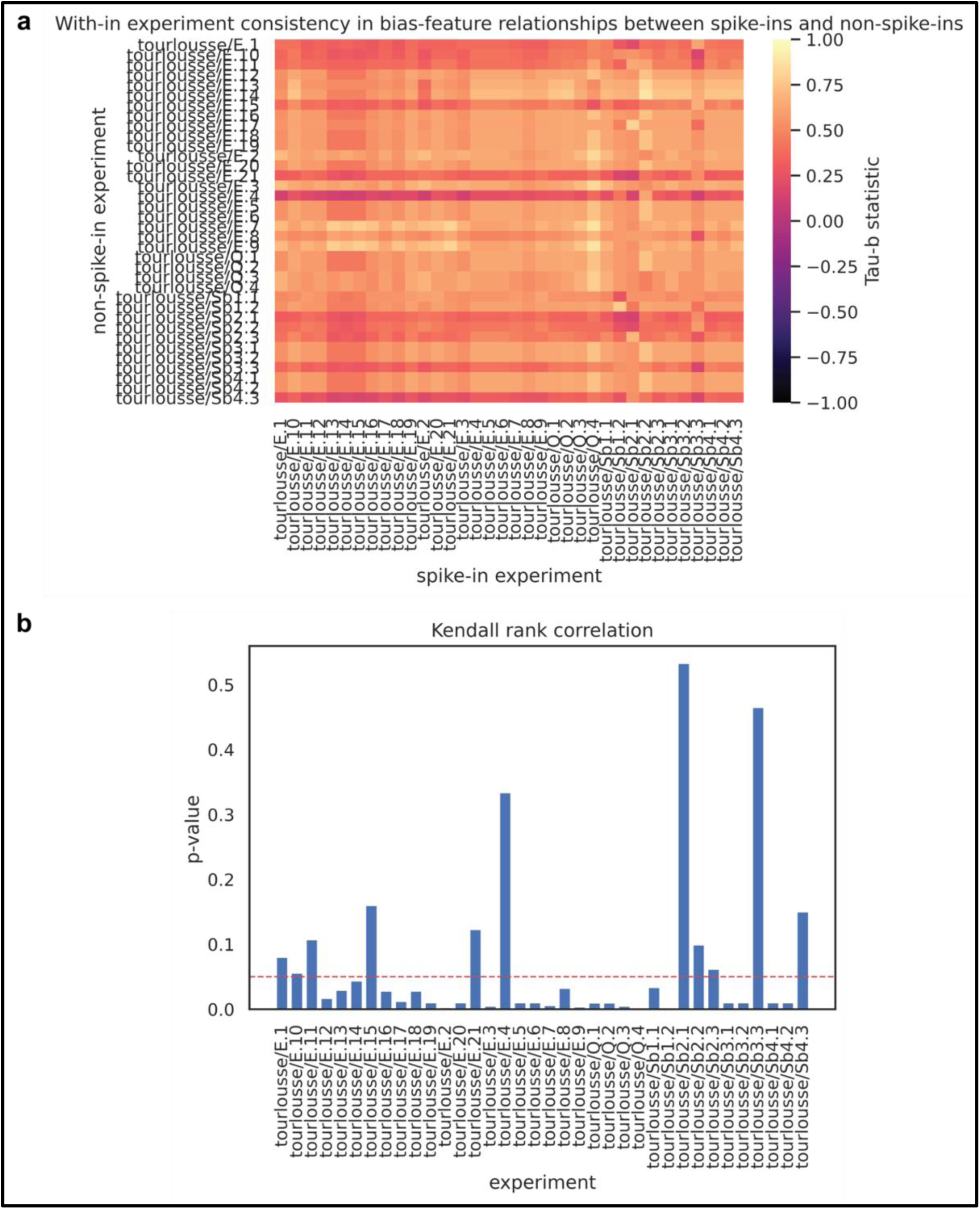
Ordinal association between SP and NSP ranked features based on bias predictability. **a** Heatmap of τ_B_ statistic between SP and NSP ranked features from various experiments of the spike-in dataset. The τ_B_ statistic was computed in a pairwise fashion between each NSP and SP experiment. Here SP/NSP *experiment_i_*refers to the SP/NSP samples within *experiment_i_* respectively. **b** P-values from the 2-tailed test under the null hypothesis that the τ_B_ statistic has an expected value of 0. The significance test was performed for SP and NSP ranked features within the same experiment, i.e. the diagonal of the heatmap in **a**.

With the findings from preliminary analysis of bias and its relationship with each *feature_i_*, we were interested in whether bias correction is a predictable task in general. Hence, we randomly split up the samples at hand into training and testing sets nondiscriminatory of dataset or experiment, such that 70% of all samples were used as training samples while the other 30% of the samples went into testing the performance of the model. By comparing the performance of each regression model to the baseline (*r_raw_*) established by the vanilla control model, we observed that modeling generally improved the error in inferring the true abundance *r_true_* of a sample (Fig. S2a). This suggested that bias correction in metabarcoding studies is a predictable task. In addition, we found XgBoost and random forest to be the two optimally performing models across the board. When further exploring the importance of each *feature_i_* in training these two models, we found that *amp_gc* stood out as a strong feature predictor consistent with the findings from the analysis of NMI (Fig. S2g, h).

Furthermore, we were interested in how much information needed to be provided to the model regarding a specific experiment *a priori* to make correction of bias in that experiment a predictable task. Stratification describes the process in which the data is divided into subgroups of training and validation sets when performing cross-validation on a given model. We performed leave-one-out cross-validation (LOOCV) with various granularities of stratification to evaluate model performance under the scenarios described in Fig. 4a. Under the dataset-specific scenario, LOOCV was performed using dataset as unit. The model was trained on the global dataset while leaving one of the datasets out, on which the validation is performed. Under the experiment-specific scenario, LOOCV was performed using experiment as unit. Only the spike-in dataset was used in the training of the model while leaving one of the spike-in experiments out for validation. The hybrid LOOCV combines both dataset-specific and experiment-specific frameworks by training on both the global and spike-in datasets and using one of the spike-in experiments for validation. During the training process for LOOCV, the hyperparameters for all models were tuned using a grid search optimization algorithm (Fig. S3). We observed model performance in LOOCV generally benefited from reducing the granularity of stratification, likely because there are more experimental conditions shared between experiments of the same dataset than those across datasets (Fig. 4b). In addition, the performance of the hybrid models appeared to be worse than that of experiment-specific models, suggesting that samples from other datasets were indirectly introducing noises into the training data that overshadowed the experiment-specific relationships between feature predictors and true labels (Fig. S4). Consistent with findings from the random train-test split evaluation, we found XgBoost and random forest to be the two optimally performing models under scenarios where experiment-specific information is provided. Under these scenarios, we also found that *amp_gc* emerged again as a strong feature predictor for these two models when evaluating the importance of each *feature_i_* (Fig. 5d-i).

**Figure 4:**
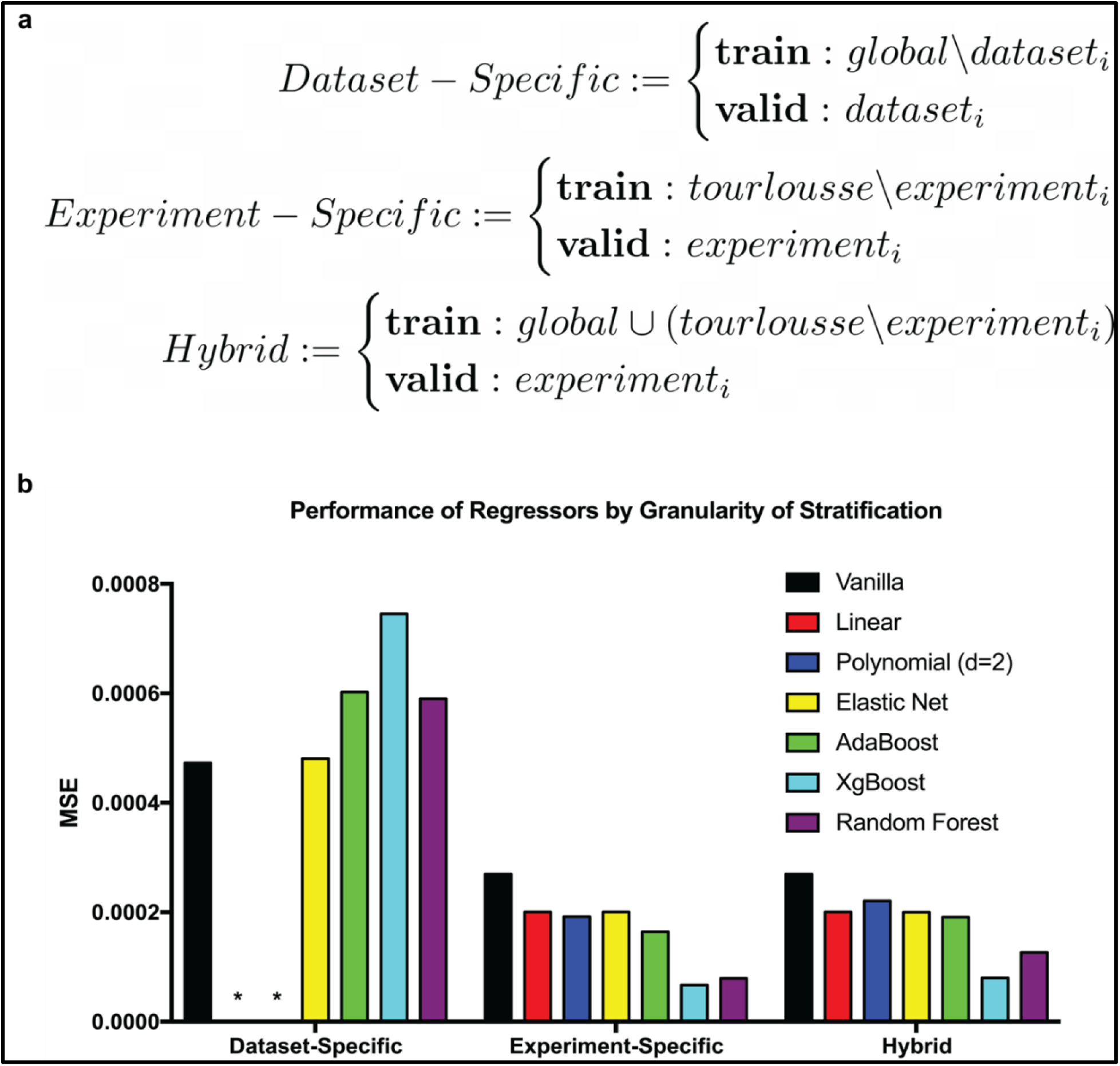
Effects of granularity of stratification on model performance. **a** Schemes of LOOCV setup and design under different scenarios. Here, *global* refers to the global dataset while *tourlousse* refers to the spike-in dataset provided by Tourlousse et al. (2017). **b** LOOCV Performance of regression models under different stratification scenarios measured by MSE. Dataset-specific linear and polynomial models performed extremely worse compared to other models due to overfitting, thus were not plotted to retain the scale of the y-axis.

**Figure 5:**
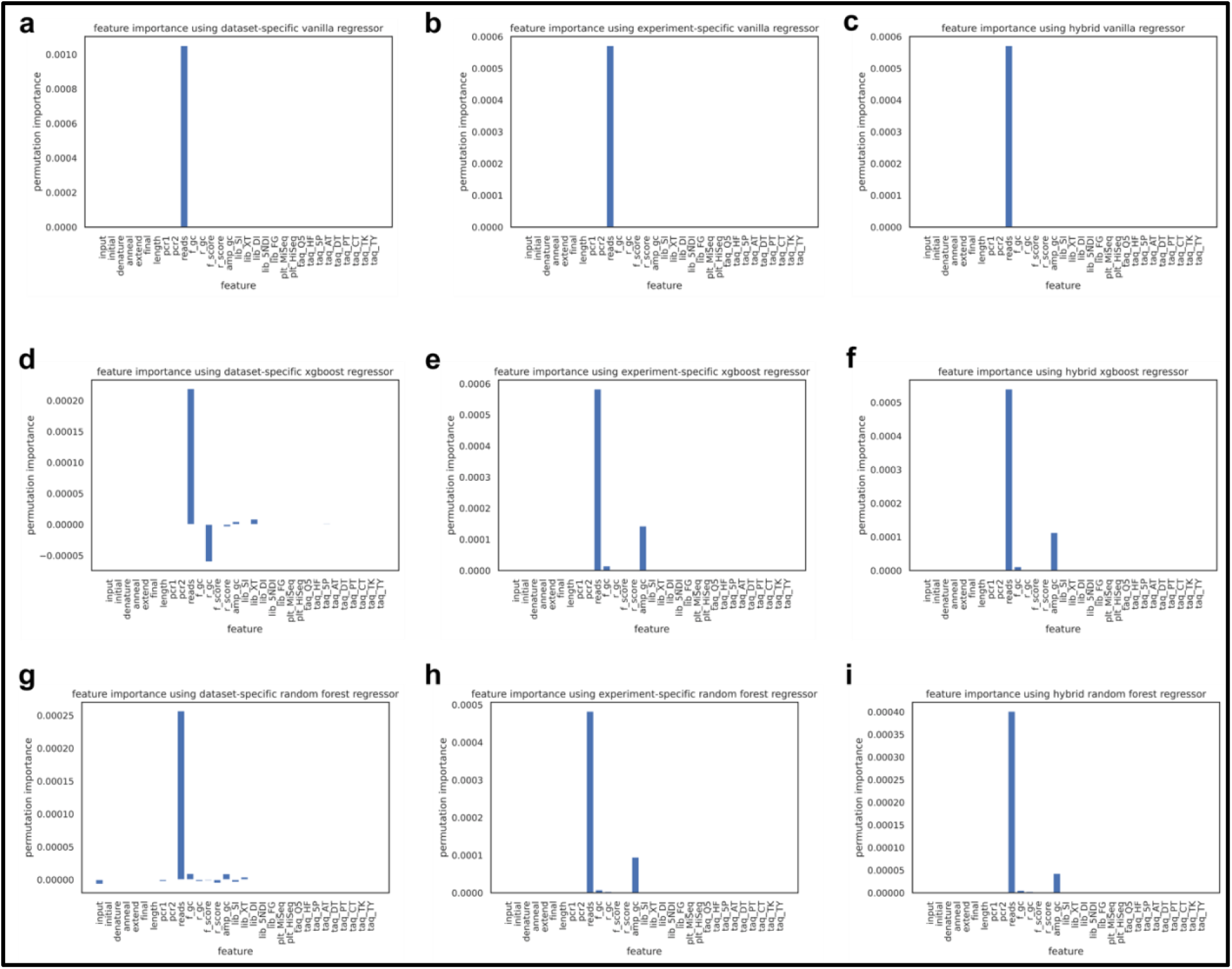
Importance of each feature_i_ in model performance under different stratification scenarios. LOOCV were performed following the schemes as described in Fig. 4a. Feature importances were measured by the permutation feature importance (PFI) algorithm developed by Fisher et al. (2018).

The consistency in feature importances prompted us to question whether only two feature predictors would suffice to achieve the same model performance. We tested this hypothesis by repeating the experiment-specific scenario following the schemes described in Fig. 4a with only two-dimensional data consisting of *reads* and *amp_gc*. We observed that most of the trends between model performances were preserved while the orderings of XgBoost and random forest were inverted. We suspected that this was due to the fact that XgBoost seemed to draw more information from features other than the two most important ones than random forest according to Fig. 5e and h.

With the optimal model and important features identified, we proceeded to evaluate the realistic applications of synthetic spike-in constructs alongside supervised learning algorithms. In a practical scenario, experiment-specific information is likely not available at the time of the experiment. Hence, researchers would realistically only have access to global dataset *a priori,* which is equivalent to the dataset-specific scenario in Fig. 4a. We evaluated the model performances with SP samples incorporated in both non-discriminative and discriminative manners (Fig. 6a). Under the non-discriminative approach, SP samples were treated indiscriminately from the other global samples by the model. On the other hand, the discriminative approach applied weighted loss function that enforces the model to be penalized more for incorrection predictions on SP samples than those on the other global samples (Fig. S6). The results from this experiment were similar to the performances of dataset-specific models in Fig. 4b in that most of the models overshot the baseline performance established by the vanilla control model except for AdaBoost, which performed 35.62% better than the baseline under both non-discriminative and discriminative approaches (Fig. 6b). The weighted loss function was only able to marginally rescue the performance of elastic net regressor to 16.87% better than the baseline. In addition to the weighted loss function, we have also attempted to incorporate SP samples under the discriminative approach via residual coefficients, which also failed to improve the baseline performance (Fig. S7a). The residual coefficient method is a procedure applied on models with linear regression function, such that a regressor is trained on global samples first while few parameters of the linear regression function are further optimized by sequentially training on the SP samples (Fig. S7b-d). The findings from Fig. 6 were similar to those from the granularity of stratification experiment in that models tend to perform poorly when experiment-specific information is not available *a priori*.

**Figure 6:**
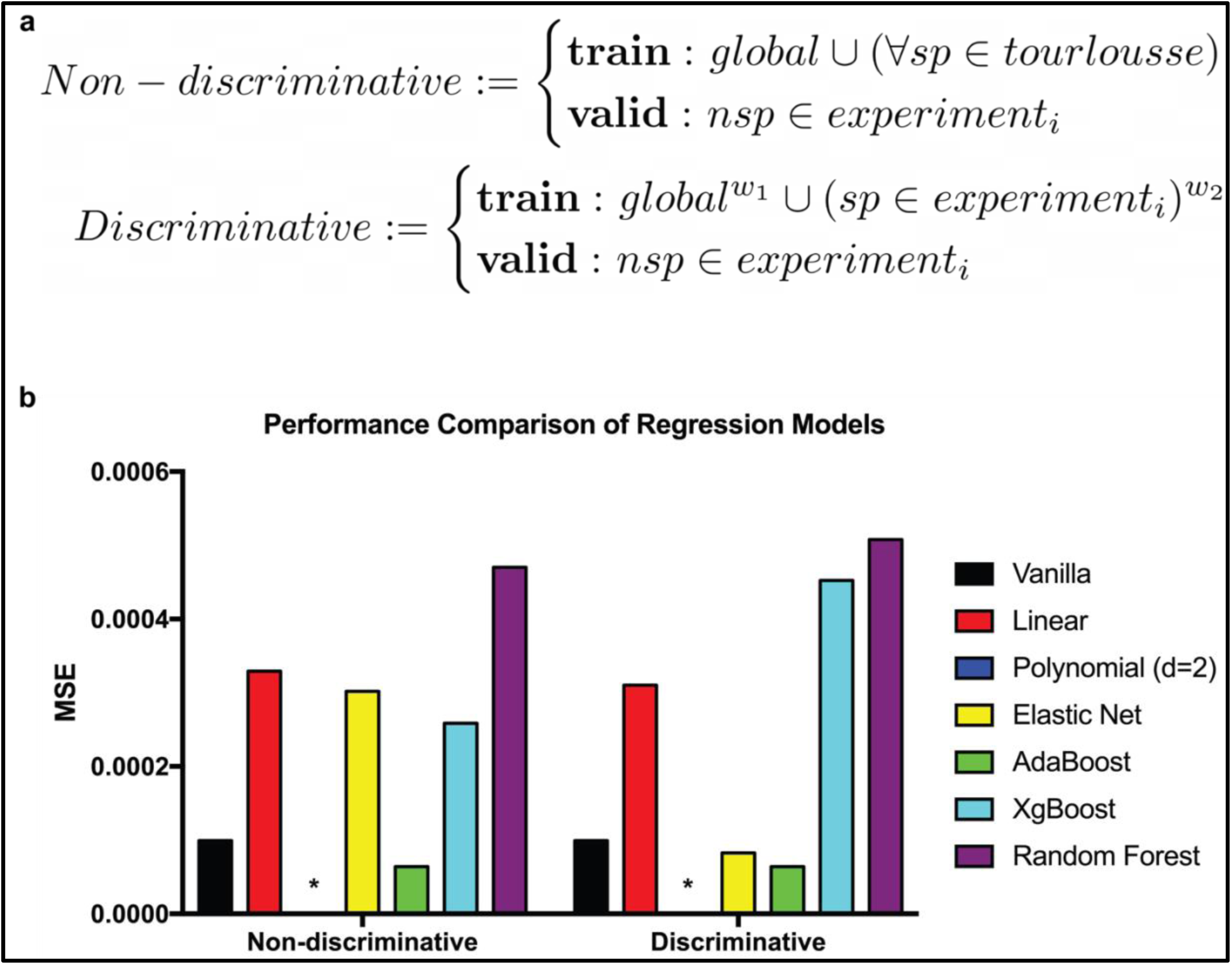
Performances of dataset-specific models with synthetic spike-in constructs. **a** Schemes of LOOCV setup and design under different spike-in integration approaches. Here, non-discriminative approach treated SP samples indiscriminately from the other global samples while discriminative approach placed an emphasis on correction predictions of SP samples over those of the other global samples. **b** LOOCV performances of dataset-specific regression models under different spike-in integration approaches measured by MSE. Polynomial models were not shown in the figure as they exhibited drastic overfitting as compared to the other models.

Drawn from the conclusions in the granularity of stratification experiment, we attempted to rescue the model performances in Fig. 6 by providing the models with experiment-specific data in the training stage (Fig. 7a). We observed that performances of the models were indeed reverted back to those observed under the hybrid scenario in Fig. 4b with XgBoost and random forest being the optimally performing models across the board when experiment-specific information is available (Fig. 7b). Specifically, XgBoost showed an optimal 81.57% improvement from the baseline under the non-discriminative approach while random forest exhibited a suboptimal 75.04% improvement from the baseline under the discriminative approach. Moreover, XgBoost was doing so without any emphasis on the presence of SP samples. The discriminative approach improved the performances of AdaBoost and random forest but did not ultimately achieve the optimal performance by XgBoost. The findings were consistent with those from the granularity of stratification experiment despite the presence of SP samples, which directed at the limitations of synthetic spike-in constructs under the current integration approaches.

**Figure 7:**
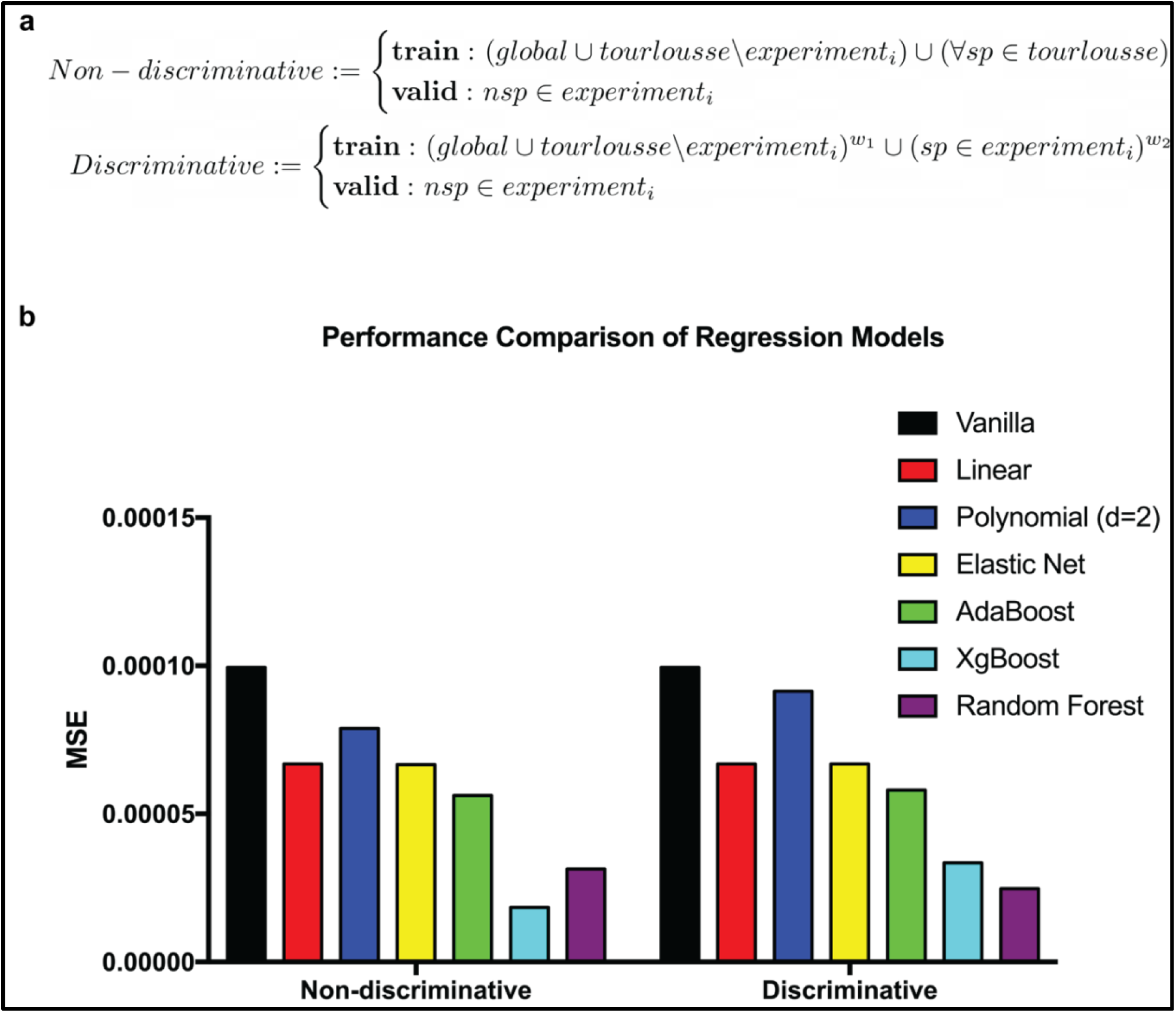
Rescued performances of hybrid models with synthetic spike-in constructs. **a** Schemes of LOOCV setup and design under different spike-in integration approaches. The definitions of non-discriminative and discriminative approaches were as described in Fig. 6a. **b** LOOCV performances of hybrid regression models under different spike-in integration approaches measured by MSE.

## Discussion

In this study, we evaluated the combined performance of SL algorithms and synthetic spike-in constructs in correcting the technical bias of eDNA metabarcoding studies. We have mined mock community datasets from nine relevant studies to simulate realistic environmental samples (Angel et al., 2018; Bokulich et al., 2015; Callahan et al., 2016; Gohl et al., 2016; Kozich et al., 2013; Leray and Knowlton, 2017; Schirmer et al., 2015; Taylor et al., 2016; Tourlousse et al., 2017). These mock communities were defined cultures of microorganisms generated *in vitro* to simulate composition of microbial communities found in nature (Highlander, 2013). In the process of mining these datasets, we were provided with raw NGS reads and various experimental conditions detailed in each study. The taxonomy of raw NGS reads were naively assigned using the taxonomic assignment package *Kraken2* to simulate presumably biased relative abundances from realistic metabarcoding workflow (Wood et al., 2019). The various experimental conditions were extracted and preprocessed as feature predictors for the downstream model construction. These included the temperature at various stages of PCR, sequenced read length, primer GC-contents, total amount of input DNA within each experiment, library preparation method, sequencing platform, and Taq polymerase. The interactions between primers and templates were analyzed by performing *in silico* PCR using the ecoPCR pipeline (Ficetola et al., 2010). Categorical features were converted to quantitative information through one-hot encoding. In addition, a primer binding score was computed for every pairwise primer-template interaction by fitting a linear model on data provided by Bru et al. in a previous study, looking at the detrimental effects of position-wise primer-template mismatches on efficiency of PCR amplification (2008).

Prior to the model construction, we looked into whether there existed information regarding the bias to be extracted from each *feature_i_* as preliminary analysis. We found that outside of *reads*, which was inherently involved in the derivation of bias, *amp_gc* emerged as another potentially strong feature predictor. In addition, we were interested in whether the ordering in strengths of feature predictors would transfer between SP and NSP samples within the same experiment. Thus, we first plotted bias against each *feature_i_* for SP and NSP samples individually and evaluated the strength of each feature predictor by calculating the MSE of a linear regression model when fitted onto the bias- *feature_i_* plot, which indicated the goodness of fit. Features were ranked based on the MSE of linear regression model for each of SP and NSP samples respectively within each experiment. We then computed the Kendall rank correlation coefficient between the SP and NSP ranked feature vectors to measure their ordinal associations. We found that the pattern in strengths of feature predictors only transferred between SP and NSP samples for a fraction of the experiments in the spike-in dataset, which hinted at the limitations of synthetic spike-in constructs in bias correction.

With potential candidates of strong feature predictors in mind, we explored whether bias correction is indeed a predictable task by randomly assigning 70% of all samples into training set and the other 30% into testing set nondiscriminatory of dataset or experiment. We found that modeling generally improved in performance compared to the baseline established by a vanilla control model. This suggested that bias correction is indeed a predictable task; upon further investigation of feature importances, we found that *amp_gc* stood out again as a strong feature predictor.

Next, we explored how much information needs to be shared between training and validation data for bias correction to be a predictable task. To that end, we performed LOOCV under three different scenarios and observed that experiment-specific information was necessary for regression models to perform well in bias correction, where XgBoost and random forest were the optimally performing models. We also found that the addition of samples from other datasets to the experiment-specific training data worsened the model performances in general, suggesting that non-experiment-specific samples could be introducing noises that contaminated the experiment-specific relationships between feature predictors and true labels. Given the consistency in *reads* and *amp_gc* being the strongest feature predictors among the rest, we tested how well each model could perform if it was only given two feature predictors to begin with. This was done by replicating the experiment-specific scenario from the granularity of stratification experiment with a two-dimensional input data containing only *reads* and *amp_gc* as features. We observed that the ranking in performance among the models were mostly preserved with XgBoost and random forest inverted. We suspected that this was likely due to XgBoost drawing more information from the other excluded features than random forest as indicated by prior analysis of feature importances.

With the promising findings in modeling for bias correction, we were interested in incorporating synthetic spike-in constructs into the existing model construction. Conventionally, spike-in controls are used to construct a normalization function that maps from r_raw_ to r_norm_ by comparing the normalized spike-in concentrations to the raw spike-in read counts in each experiment, thus standardizing the read counts across multiple experiments of the same study (Chen et al., 2016). With SL algorithms, we could devise a flexible approach that bypasses the batch-wise normalization step altogether. This could be achieved by constructing a robust enough model that is able to take in both experimental conditions and *r_raw_* as feature predictors to infer *r_true_* directly.

In a realistic application of spike-in-mediated bias correction, it is less likely that any experiment-specific data would be available at the time of the experiment unless the experiment is replicating an experiment from prior study. Hence, we first evaluated the performances of various models under the dataset-specific scenario with SP samples integrated. SP samples were integrated in two fashions, non-discriminative and discriminative approaches. The non-discriminative approach did not differentiate SP samples from any other global samples while the discriminative approach used weighted loss function to penalize the model more for incorrect predictions on SP samples than those on other global samples. Under the dataset-specific scenario, we found AdaBoost to be the optimally performing model with both non-discriminative and discriminative approaches, yielding a 35.62% improvement from the baseline performance established by the vanilla control model. Aside, we only observed marginal improvements from the discriminative approach compared to the non-discriminative approach.

Given the results from the granularity of stratification experiment, we suspected the integration of SP samples into dataset-specific models primarily failed due to the lack of experiment-specific information to begin with. Accordingly, we attempted to rescue the model performances by replicating the SP integration with hybrid models from the granularity of stratification experiment instead. As expected, the model performances were reverted back to those observed under the hybrid scenario in the granularity of stratification experiment. Moreover, XgBoost showed an optimal 81.57% improvement under the non-discriminative approach while random forest showed a suboptimal 75.04% improvement under the discriminative approach.

Overall, we demonstrated that bias correction in eDNA metabarcoding studies is indeed a predictable task. Likewise, future metabarcoding studies are likely to benefit from SL-based bias correction prior to downstream analyses. More specifically, our work hereby suggests that if experiment-specific is available a priori, it is optimal to start with an XgBoost model; otherwise, it is best to start with an AdaBoost model that is still a marginal improvement from the baseline with no modeling. Furthermore, there exist many questions that remain to be addressed in future studies. We found negligible improvements between the non-discriminative and discriminative approaches of SP integration, which led us to question whether weighted loss function and residual coefficient are the optimal way of incorporating SP samples into the model construction. On the same note, it could be that case that our training data did not have sufficient amount of SP samples per experiment. Currently, the experiments from the spike-in dataset each contains 12 SP samples per experiment. Lastly, we suspect that quantitative information regarding the different Taq polymerases would further improve the model performances as the current one-hot encoding approach is unable to capture the functional and structural characteristics of each Taq polymerase. Addressing each of these questions in future studies would allow one to construct better bias correcting models that potentially lead to more accurate inference of microbial community composition in eDNA metabarcoding studies.

## Conclusions

Using mock community datasets mined from nine prior studies, we showed that bias correction in eDNA metabarcoding studies is indeed a predictable task. In other words, it is possible to bypass the conventional batch-wise normalization step and infer r_true_ directly using various experimental conditions and r_raw_ as feature predictors. We found *reads* and *amp_gc* to be the two strongest feature predictors, such that the two features alone are sufficient to retain most of the model performances. Based on different stratification scenarios in LOOCV, we concluded that experiment-specific information is necessary for bias correcting models to perform well. Yet, we have not discovered an effective method of integrating the SP samples with the existing model training framework as none of our optimally performing models placed an emphasis on the presence of SP samples discriminatively. More specifically, under the data-specific scenario, AdaBoost showed an optimal 35.62% improvement from the baseline with both non-discriminative and discriminative approaches. With experiment-specific data provided, XgBoost exhibited an optimal 81.57% improvement from the baseline using the non-discriminative approach. With all findings considered, our work suggests that it would be beneficial for future metabarcoding studies to perform SL-based bias correction prior to downstream analyses. Furthermore, we have found that it is optimal to begin with an XgBoost model if experiment-specific data is available. Otherwise, it is still recommended to begin with an AdaBoost model, which performs marginally better than the baseline with no modeling.

## Acknowledgements

I am deeply grateful to Professor Rasmus Nielsen for his mentorship and guidance, which have been instrumental in my growth as an independent scientist. I also sincerely appreciate the support and insights of Ammon Corl, Tyler Linderoth, Aaron Stern, Sandra Hui, Debora Brandt, Lenore Pipes, and Lydia Smith throughout my time in the Nielsen Lab. Additionally, I extend my gratitude to Professors Rosemary Gillespie, Jonathan Shewchuk, John Kuriyan, Sofia Villas-Boas, and the Rausser College Honors Program for their contributions to this thesis. This work was supported by the Extreme Science and Engineering Discovery Environment (XSEDE) Bridges system at the Pittsburgh Supercomputing Center through allocation BIO180028.

## Author Contributions

JC drafted the manuscript. JC and RN conceptualized the methods and experimental designs while JC performed the experiments and analyses on the methods. RN provided supervision throughout the entire workflow of the investigation. Both authors provided feedbacks regarding revisions of the manuscript, read, and approved the final manuscript.

## Availability of data and materials

The datasets and models for replicating the findings and conclusions of this article are included in the additional and supplementary information sections of this article.

## Competing interests

The authors declare that they have no competing interests.

## Supplementary Materials

### Materials and Methods

#### Feature engineering

##### Taxonomic assignment

Taxonomy of the sequenced eDNA reads from the mock community datasets were assigned using a vanilla taxonomic assignment pipeline, *Kraken2*, to mimic presumably biased relative abundances among taxa from realistic experiments. *Kraken2* was selected among many other taxonomic assignment packages for its time and memory efficiency without sacrificing much of the accuracy in assignments (Wood et al., 2019). Synthetic spike-in constructs were manually added to the standard *Kraken2* database as pseudospecies with unique taxonomic identifiers.

##### *In silico* PCR

*In silico* PCR of each universal primer pairs was performed against NCBI reference sequence database using the package, *ecoPCR*, to capture amplification dynamics of all potential amplicons (Ficetola et al., 2010). Synthetic spike-in constructs were manually added to the NCBI reference sequence database as pseudospecies with unique taxonomic identifiers. For the analyses presented in this article, we considered up to maximum of three mismatches between each primer and the template to be a successful amplification with amplicon length under 2kbp.

##### Primer binding score

A linear model was constructed to estimate the position-wise detrimental effect of a single primer-template mismatch using experimental data collected by Bru et al. in a prior study, quantifying the effects of primer-template mismatches on efficiency of PCR amplification of the 16S rRNA gene (2008). The 16S rRNA gene copy numbers from *Pseudomonas aeruginosa PAO1, Agrobacterium tumefaciens C58, and Sinorhizobium meliloti 1021,* were aggregated by averaging at each mismatch location first before fitting of the generalized linear model.

#### Data preprocessing

##### One-hot encoding

Categorical features, such as library preparation methods, sequencing platforms, and Taq polymerases, were converted into quantitative features using one-hot encoding defined as follows:

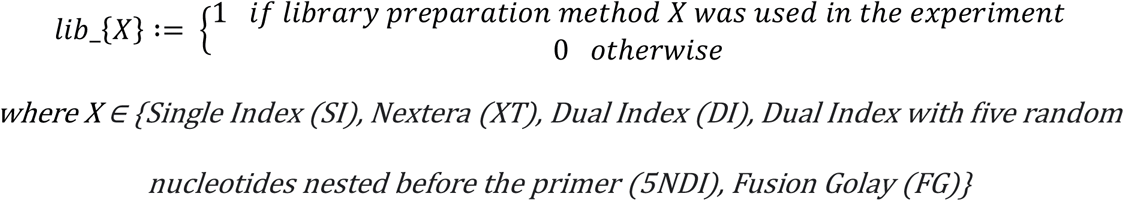

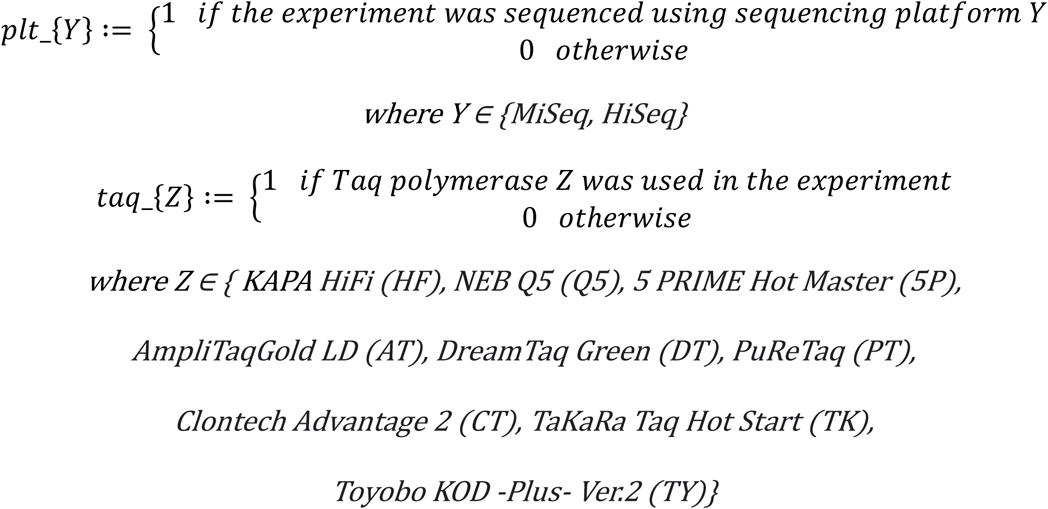

##### Read count normalization

Raw read counts were converted into relative abundances by dividing the *r_raw_* and *r_true_* of each sample by the total observed and true read counts in each experiment respectively. This is done by applying the normalization function defined as follows:

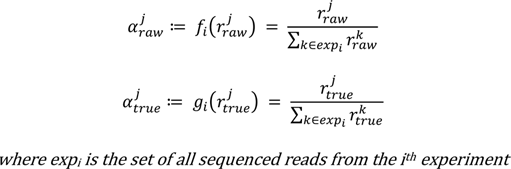

As a result, the total observed and true relative abundances must each sum up to one within each experiment.

#### Analysis of bias

For consistency, the bias of an experiment analyzed in this study is derived from Kong and Dietterich’s definition of the statistical bias of a learning algorithm as follows (1995):

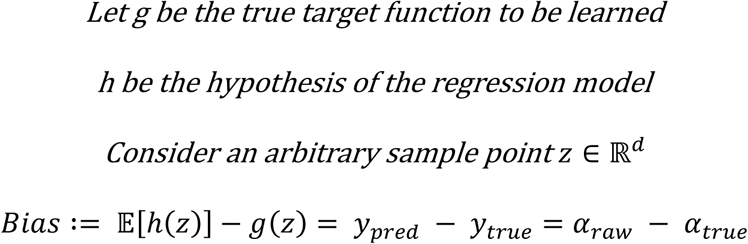

#### Mutual information

Mutual information (MI) of two random variables (RV) quantifies the amount of information obtained about distribution of one RV through observations of the other RV and vice versa. To make the values of MI comparable to other correlation coefficients, we instead used the normalized variant of MI as defined by Lancichinetti et al. as follows (2009):

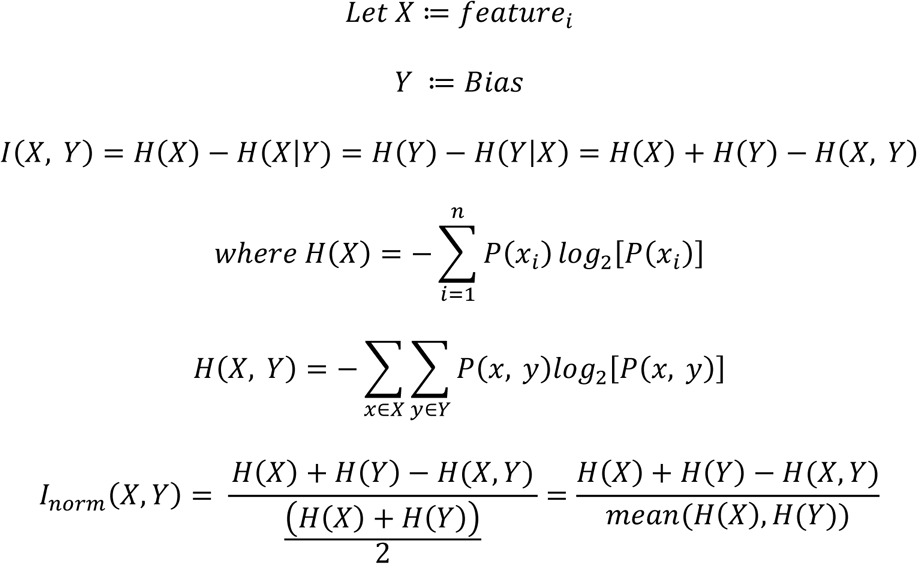

This restricts the values of MI to be in the range [0, 1] instead of [0, ∞) as in the unnormalized case.

#### Kendall rank correlation coefficient

The Kendall rank correlation coefficient, also sometimes referred to as Kendall’s τ coefficient, quantifies the ordinal association between two quantities, X and Y. In other words, Kendall’s τ coefficient measures whether ordering of one quantity reveals information about ordering of the other quantity and vice versa. In this study, the τ_B_ variant of Kendall rank correlation coefficient was adopted to account for potential ties and normalize the statistic such that a τ_B_ of -1 indicates perfect inversion while a τ_B_ of +1 signifies perfect agreement. Formally, the definition of the Kendall τ_B_ statistic is as follows (Agresti, 2010):

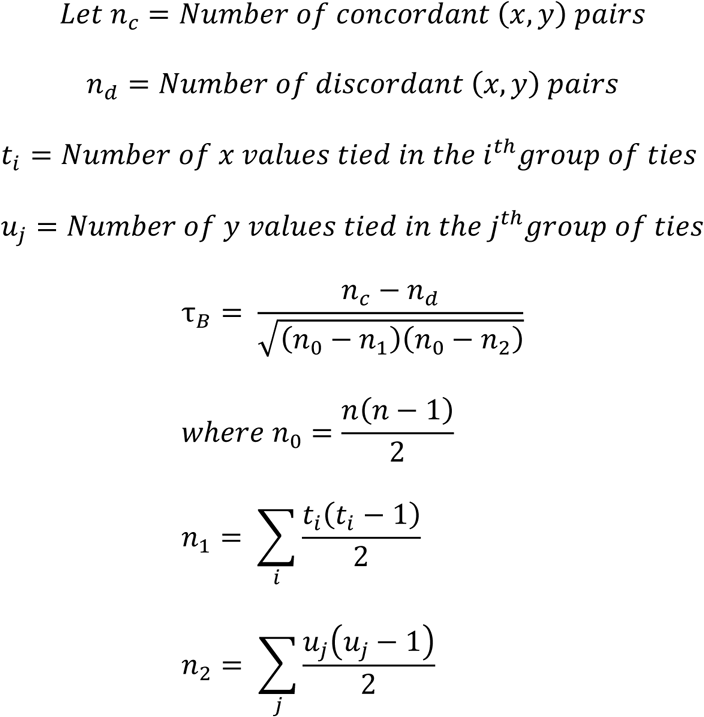

When used as a test statistic in a statistical hypothesis test, the null hypothesis for the 2-tailed test is that the expected value of τ_B_ is 0. In other words, the two quantities, X and Y, are ordinally independent of each other.

#### Supervised learning algorithms

Six supervised learning algorithms were evaluated in the present alongside an identity function as a vanilla control model, i.e. linear regression, polynomial regression with degree of 2, elastic net regression, AdaBoost, XGBoost, and random forest. Linear regressor attempts to model the distribution of object label as a linear combination of the feature predictors while degree-2 polynomial regressor includes interaction terms up to degree of 2 (Jobson, 1991). Elastic net regressor is a regularized variant of linear regressor that combines a mixture of both L1 and L2 regularizations (Zhang et al., 2017). In other words, elastic net regressor is a hybrid of lasso and ridge regressors. AdaBoost, abbreviated for Adaptive Boosting, is an ensembled regressor that trains multiple weak learners sequentially, which in this case are decision stumps with single split node, on weighted sample points (Freund and Schapire, 1997). The prediction at each time step is computed as the weighted median prediction of the weak learners currently in the ensemble, under which more accurate learners are assigned larger weights. The weight of each sample point is determined by the error of the prediction made by the weak leaners at current time step. XGBoost, short for eXtreme Gradient Boosting, is similar to AdaBoost in that it is also a boosting ensembled regressor that adds new weak learners to the ensemble sequentially. More specifically, XGBoost is a gradient boosting algorithm that is optimized for model performance and scalability (Chen and Guestrin, 2016). Under the gradient boosting method, at each iteration, a new weak learner is trained to predict the errors of the prior ensemble, such that the ensembled prediction is the sum of all weak learners in the ensemble. Gradient boosting method applies gradient descent algorithm when training a new weak learner to minimize the loss between ensembled prediction and true label. Random forest is an ensembled regressor that uses Bootstrap AGGregatING (Bagging) method, such that each weak learner is trained on a random subsample of the original training set with replacement. In addition, random forest consists of an ensemble of decision trees, in which each treenode is split on a random subset of the original features. The prediction of a random forest is computed by averaging the predictions of all decision trees in the random forest. Unless otherwise specified, all models evaluated in the present study were trained to minimize the mean squared error (MSE) between the model prediction and true label. The hyperparameters of each model were tuned using a grid search optimization algorithm where the performance of each hyperparameter combination was evaluated on the validation set (Claesen and De Moor, 2015).

#### Weighted loss function

The experiment-specific models were trained with the prior knowledge that the experiment-specific training samples more closely resemble the dynamics of the experiment than global training samples. To translate such knowledge into the model training process, we defined a specific weighted loss function, such that incorrect predictions on experiment-specific training samples are penalized more than those on global training samples, as follows:

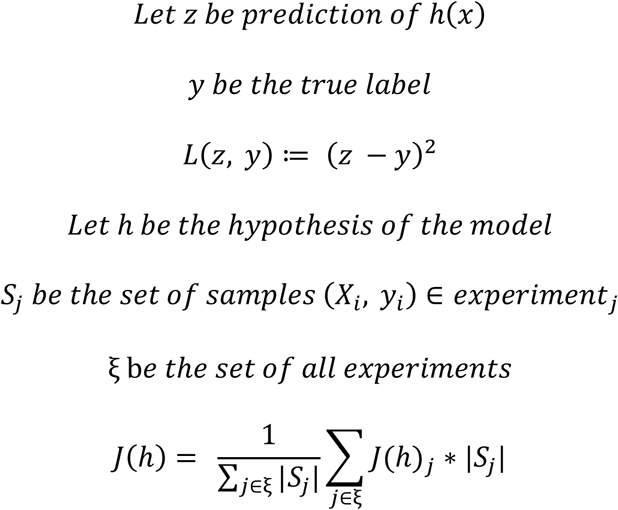

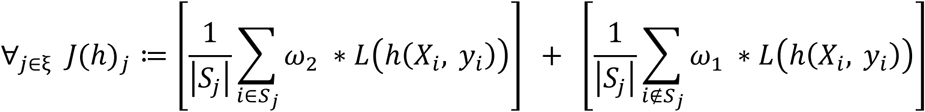

#### Feature importance

The importance of each feature within a model was measured using the permutation feature importance (PFI) algorithm as described in Fisher et al. (2018). The PFI algorithm first trains the model on the training set and predicts on the complete validation set to establish the baseline for optimal model performance. The importance of each *feature_i_* is measured by the increase in prediction error after shuffling *feature_i_*’s values in the validation set when compared to the baseline. Shuffling of *feature_i_* in the validation set essentially breaks any association between i^th^ feature and the object label of each sample as if the i^th^ feature was absent from the dataset.

### Abbreviations

eDNA: environmental DNA
PCR: polymerase chain reaction
NGS: next generation sequencing
r_raw_: raw read count
r_norm_: normalized read count
r_true_: true read count
SL: supervised learning
ML: machine learning
NMI: normalized mutual information
feature_i_: the ith feature
SP: spike-in samples
NSP: non-spike-in samples
MSE: mean squared error
LOOCV: leave-one-out cross-validation
α_raw_: raw abundance
α_true_: true abundance
Bagging: Bootstrap AGGregatING method
PFI: permutation feature importance algorithm

### Supplementary information

#### Data Preprocessing

This section provides an overview of the bioinformatics procedures and command lines performed to extract and preprocess features from the mock community datasets.

##### Kraken2

The paired sequenced reads were taxonomically assigned by *Kraken2* to simulate presumably biased relative abundances from naive metabarcoding workflows, as follows:

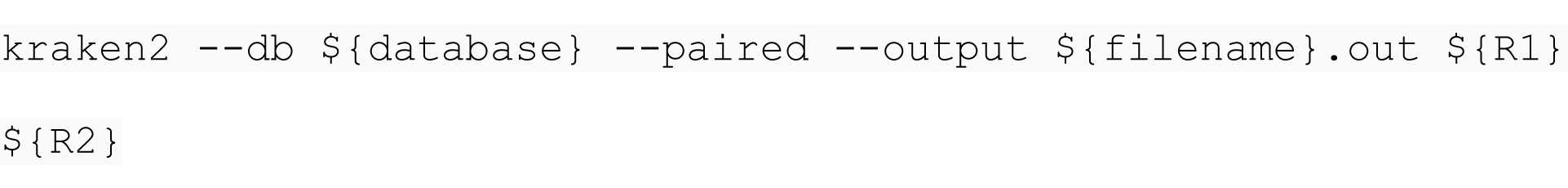

##### ecoPCR

The pairwise interactions between primers and templates were characterized by performing *in silico* PCR using the ecoPCR pipeline, as follows:

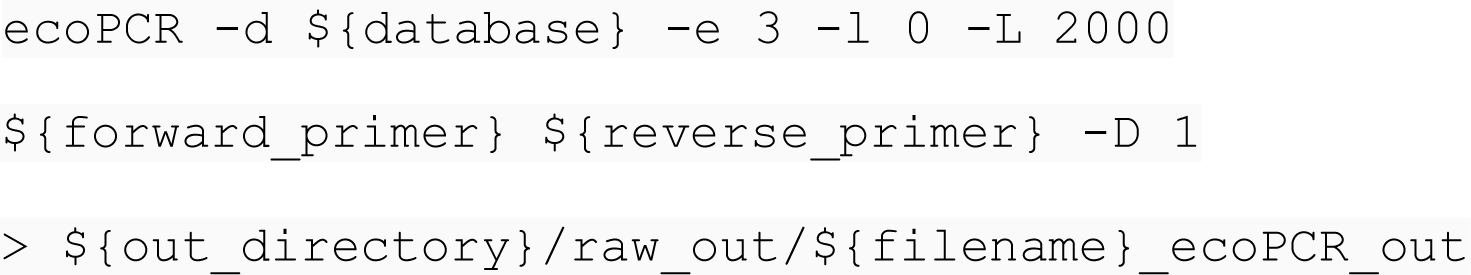

#### Residual Coefficient Method

The residual coefficient method is a procedure developed based on the hypothesis that some bias-*feature_i_* associations are conserved globally while others are experiment-specific. To translate such knowledge into the model training process, we implemented the residual coefficient method on models with linear regression function, such that a regressor is initially trained on global data to produce a set of global parameter estimates. Subsequently, few parameters of the linear regression function are tuned by sequentially training on experiment-specific SP samples. This can be illustrated formally as follows:

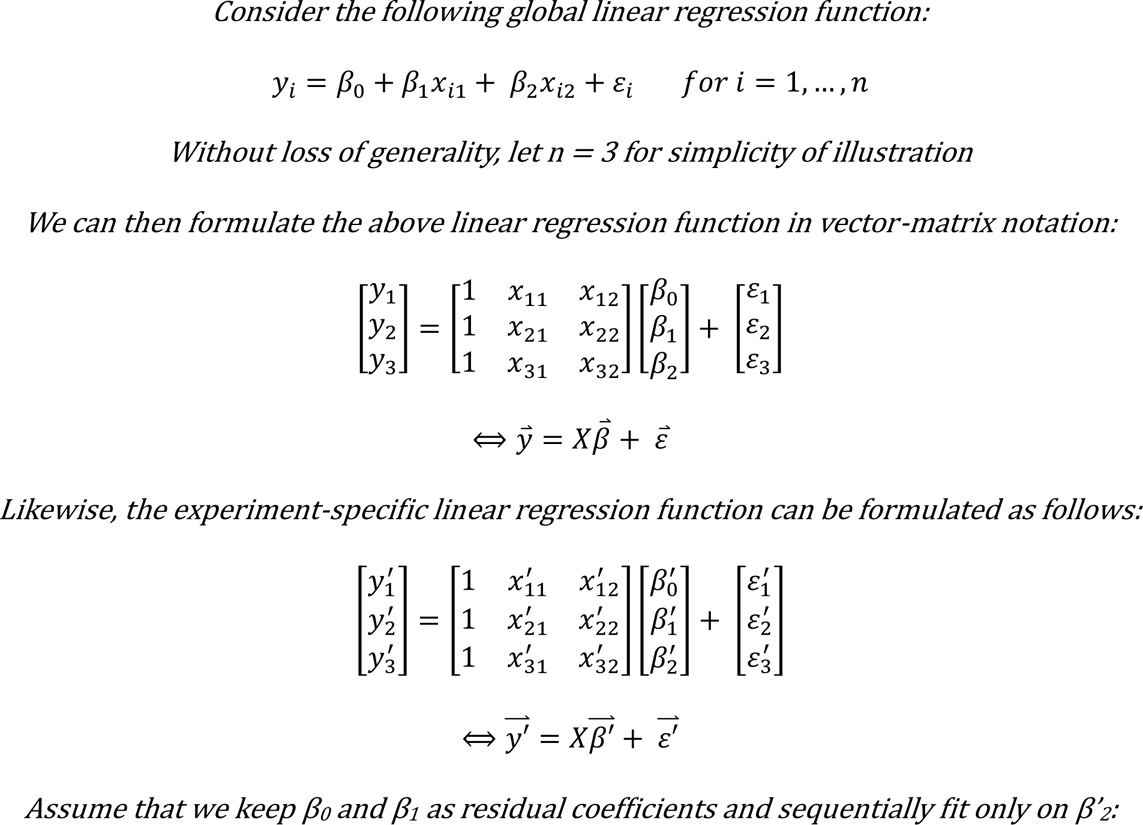

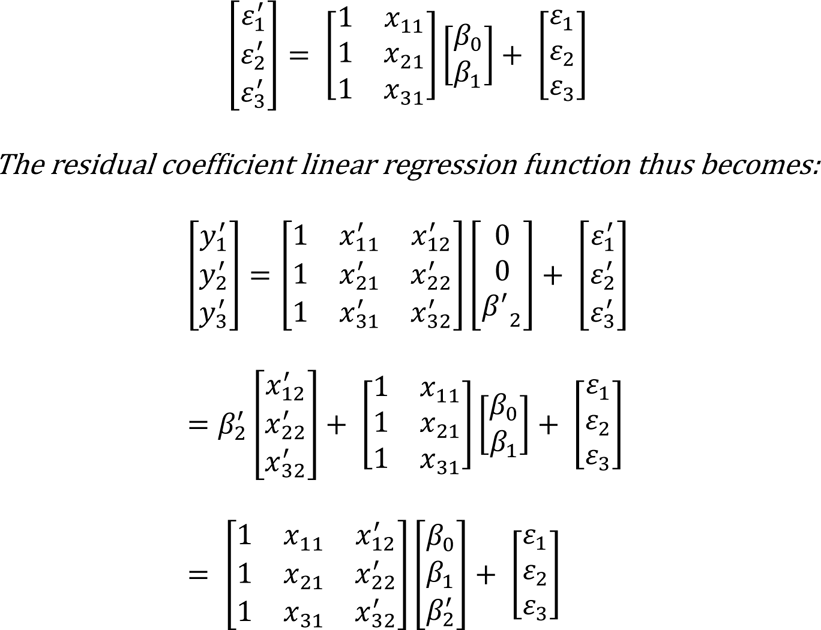

#### Optimal Models

This section provides a summary of the optimally performing bias correction models under different scenarios and approaches along with each of their tuned hyperparameters.

##### Dataset-specific model

The optimally performing AdaBoost regression model under the dataset-specific scenario with SP integration showed a 35.62% improvement from the baseline and was constructed as follows:

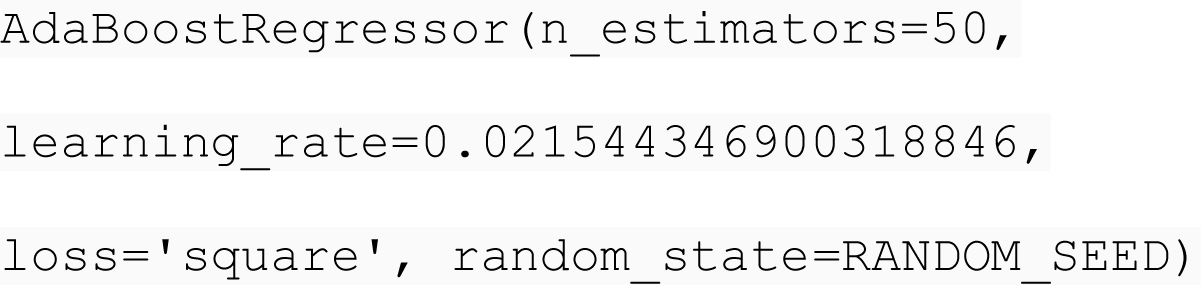

##### Rescued hybrid model

With experiment-specific data made available, the optimally rescued XgBoost regression model showed an 81.57% improvement from the baseline and was constructed as follows:

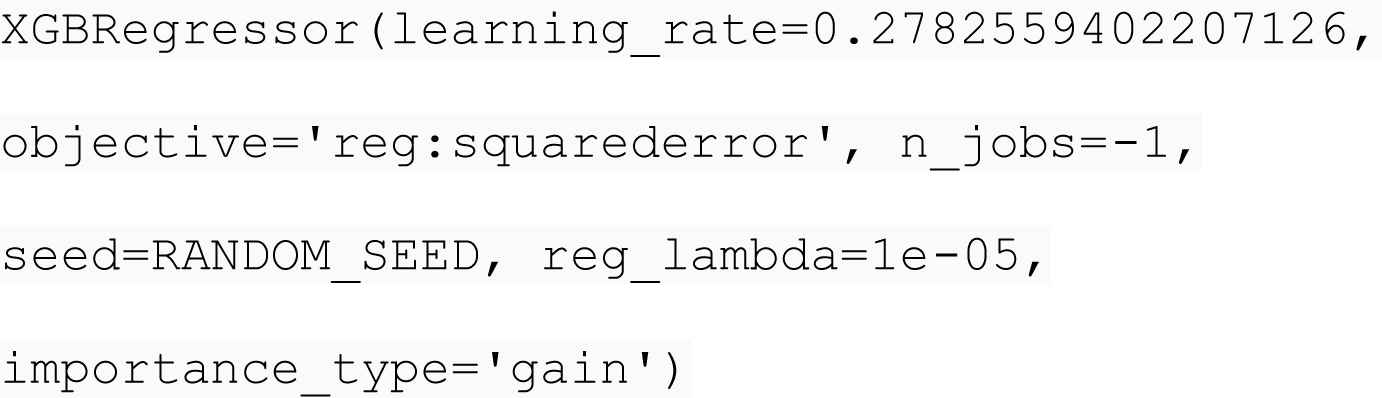

---

**Figure S1:**
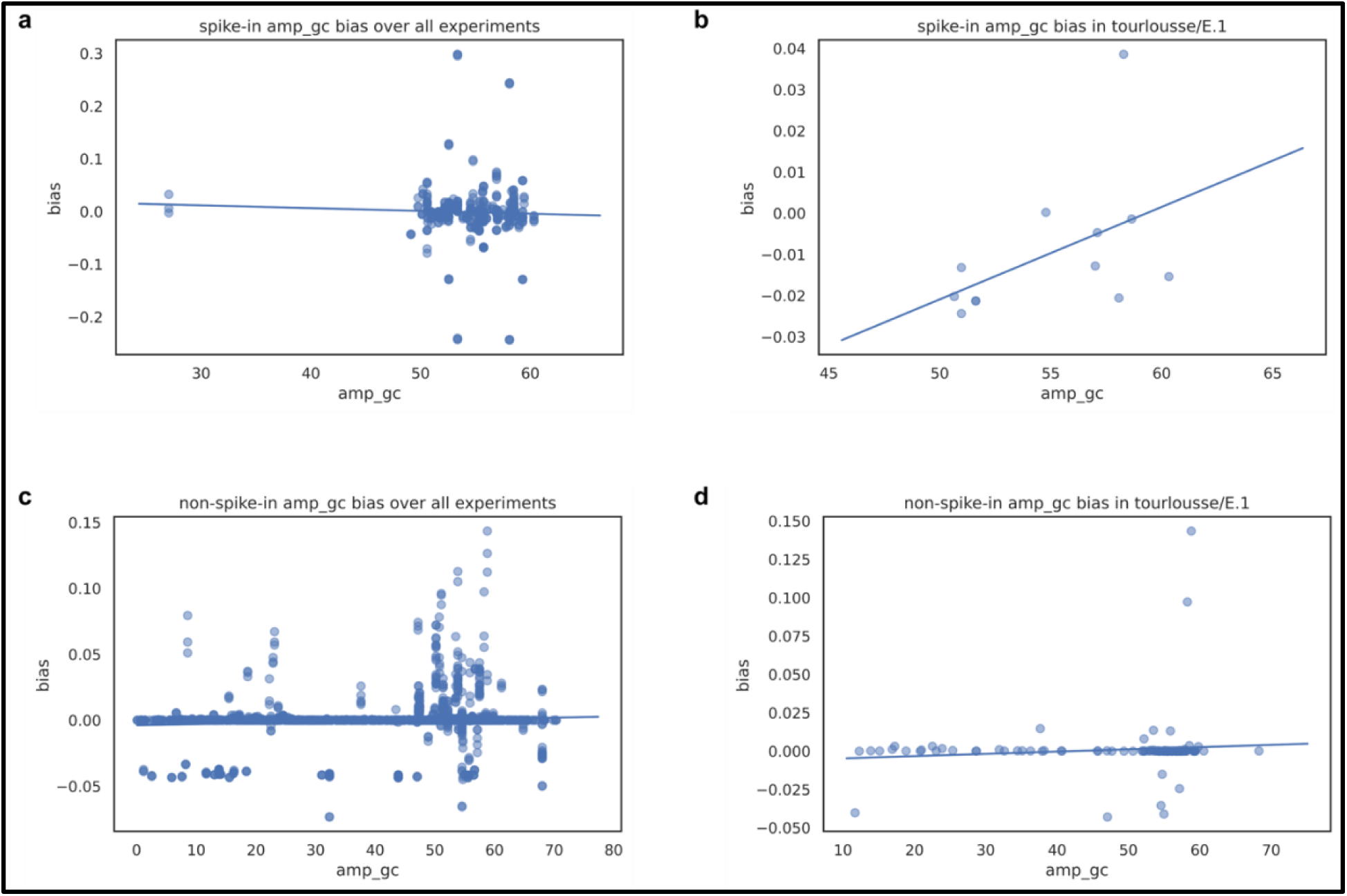
Examples of bias vs. *feature_i_* plots. To compare the SP and NSP bias-*feature_i_* associations, we plotted bias against each *feature_i_* for SP and NSP samples individually within each experiment (**b, d**) and globally across all of the experiments of the spike-in dataset (**a, c**).

**Figure S2:**
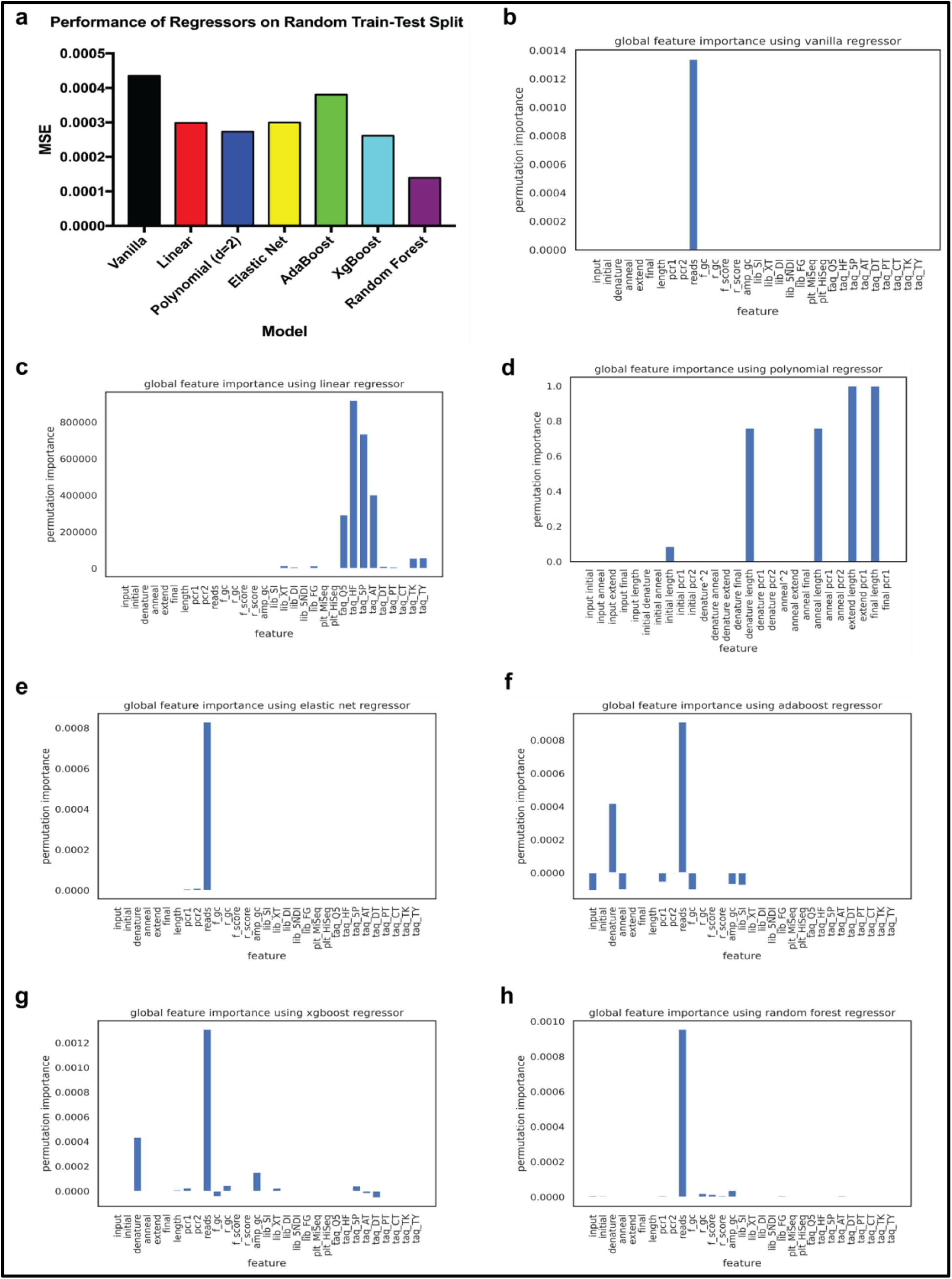
Performances of regression models with random train-test split. **a** Validation performances of regression models measured by MSE using 70% of all available samples randomly assigned as training set and the remaining 30% as testing set regardless of dataset or experiment. **b-h** Feature importances of each regression model measured by the permutation feature importance (PFI) algorithm developed by Fisher et al. (2018).

**Figure S3:**
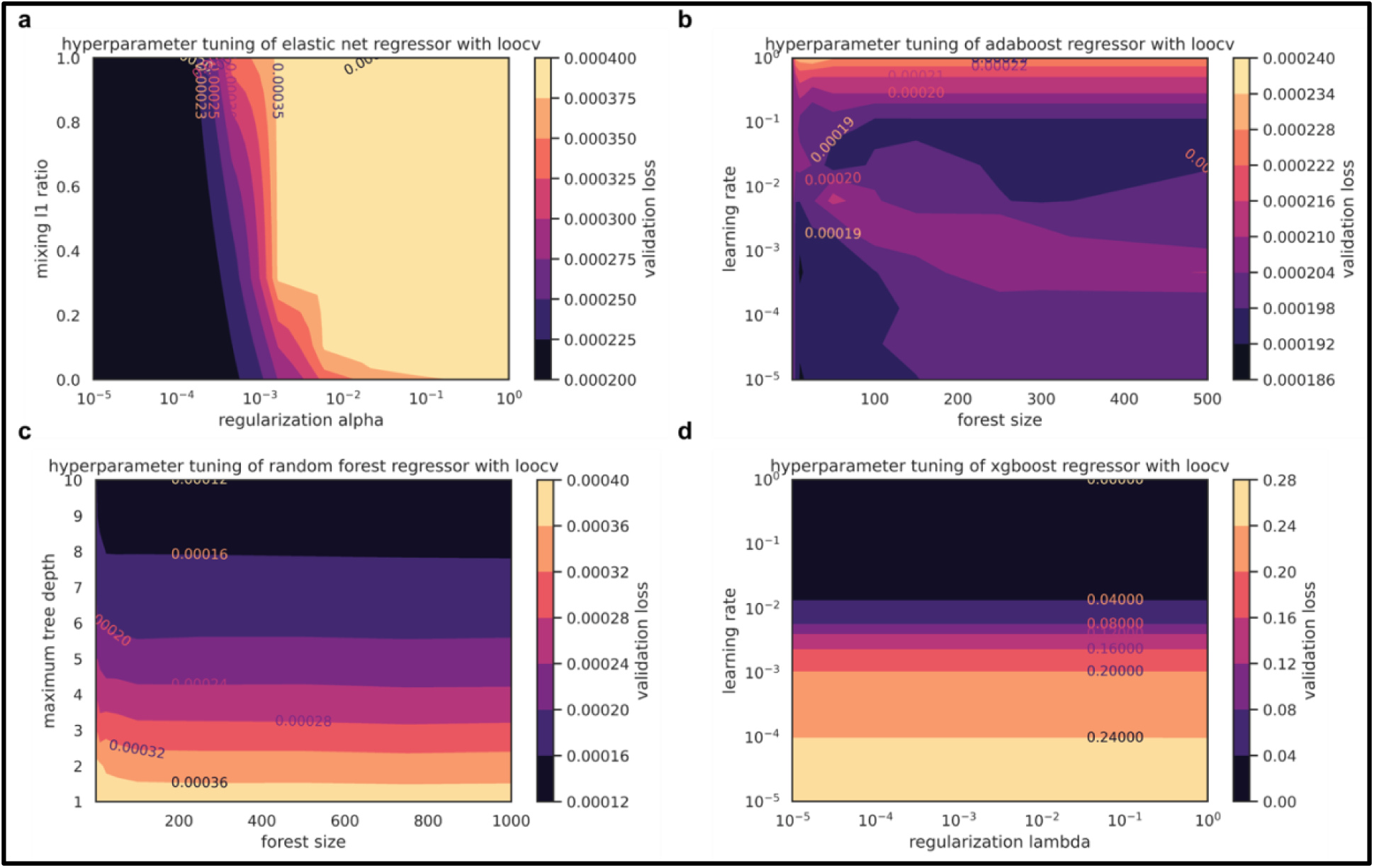
Heatmaps of hyperparameter tuning for various regression models. Hyperparameter tuning for each model were carried using a two-dimensional grid search optimization algorithm. The example heatmaps were obtained from the optimally performing experiment-specific models in the granularity of stratification experiment.

**Figure S4:**
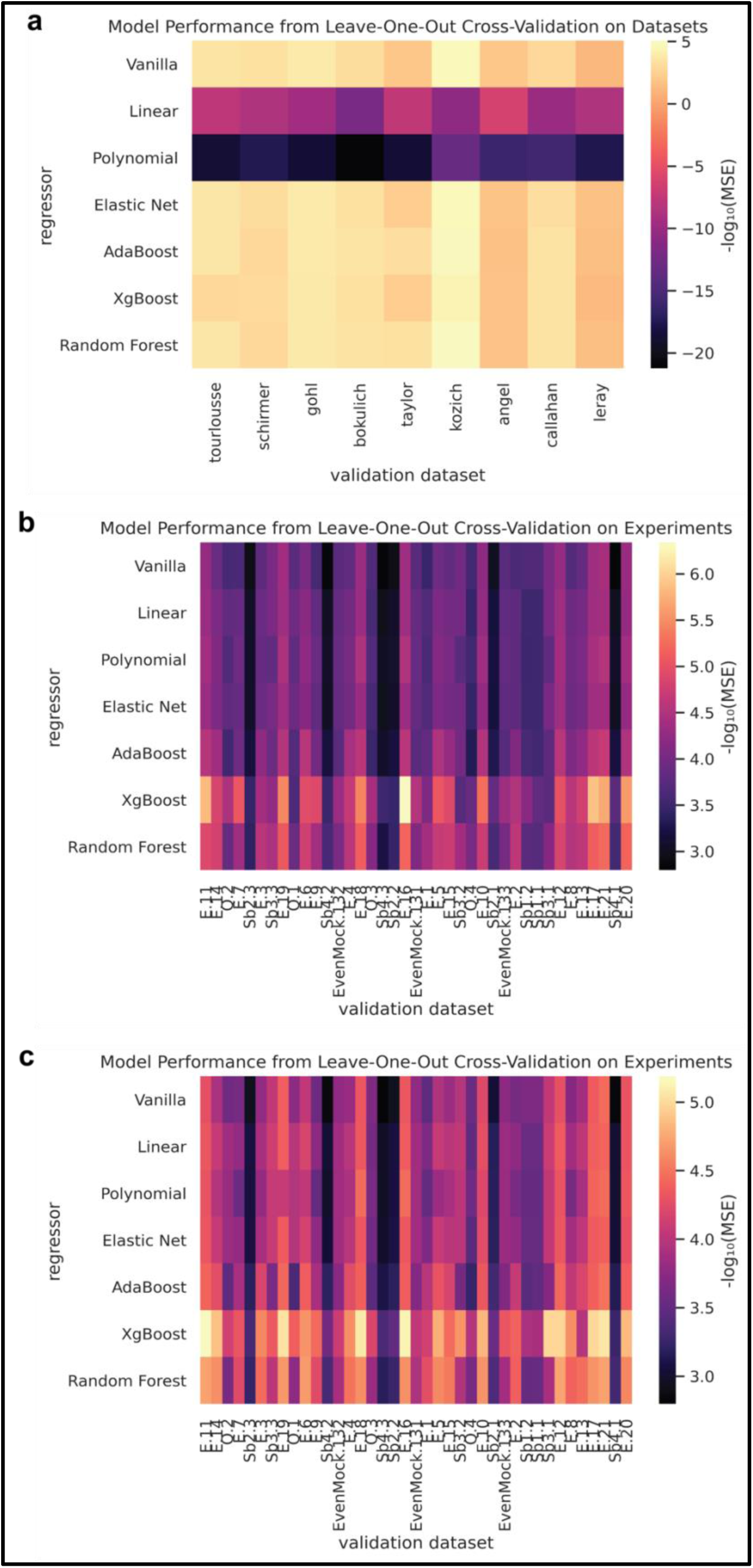
Heatmaps of model performances from the granularity of stratification experiment. Performances of regression models for each subsample of the validation set were measured by MSE under the dataset-specific (**a**), experiment-specific (**b**), or hybrid (**c**) scenario of the granularity of stratification experiment. The heatmaps were plotted in the negative logarithmic scale of the MSE for the ease of visualization.

**Figure S5:**
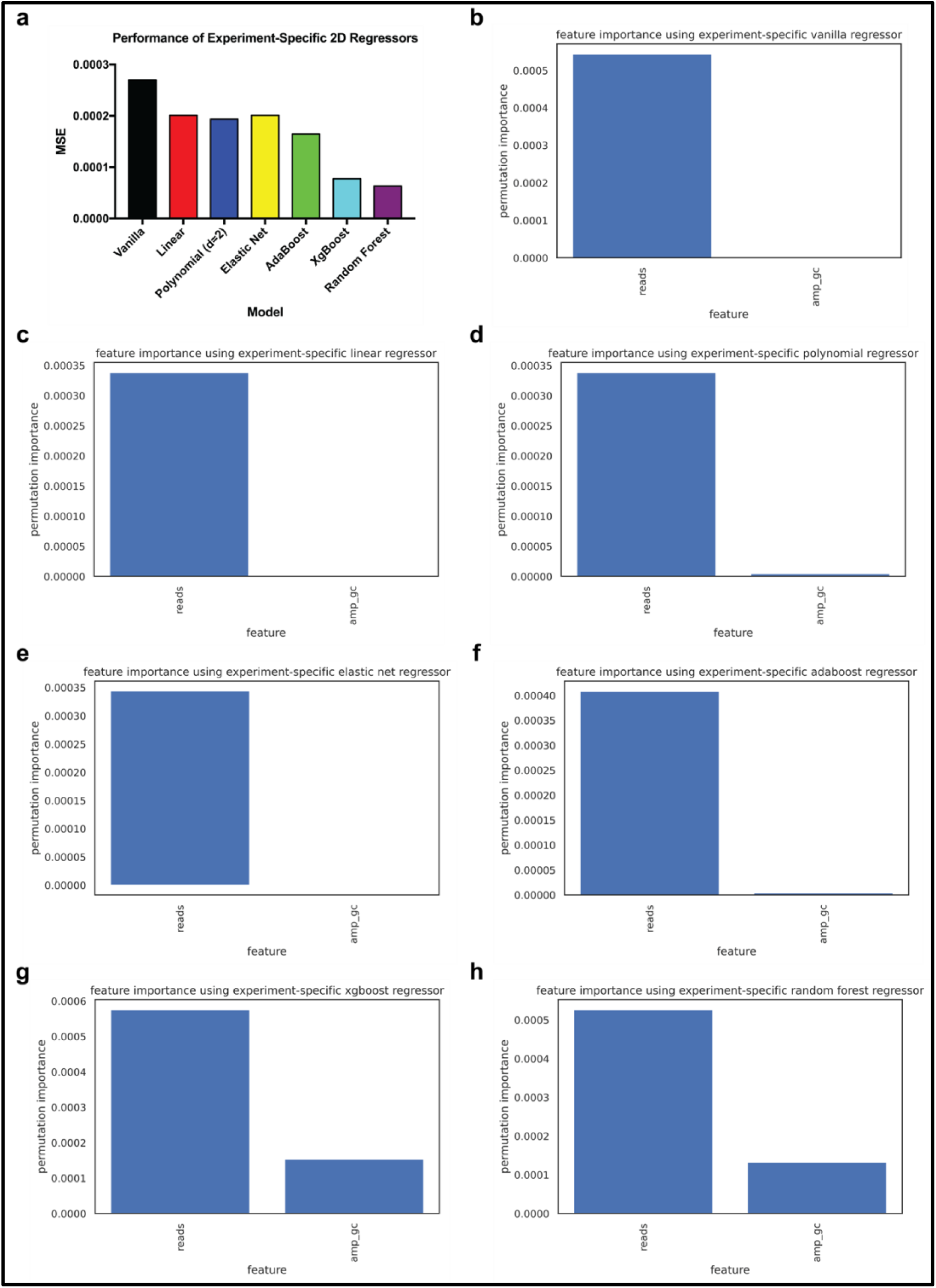
Performances of experiment-specific models with two-dimensional input data. **a** LOOCV performances of regression models following the experiment-specific scenario defined in the granularity of stratification experiment using only two feature predictors, *reads* and *amp_gc*. **b-h** Importances of *reads* and *amp_gc* in each regression model measured by the permutation feature importance (PFI) algorithm developed by Fisher et al. (2018).

**Figure S6:**
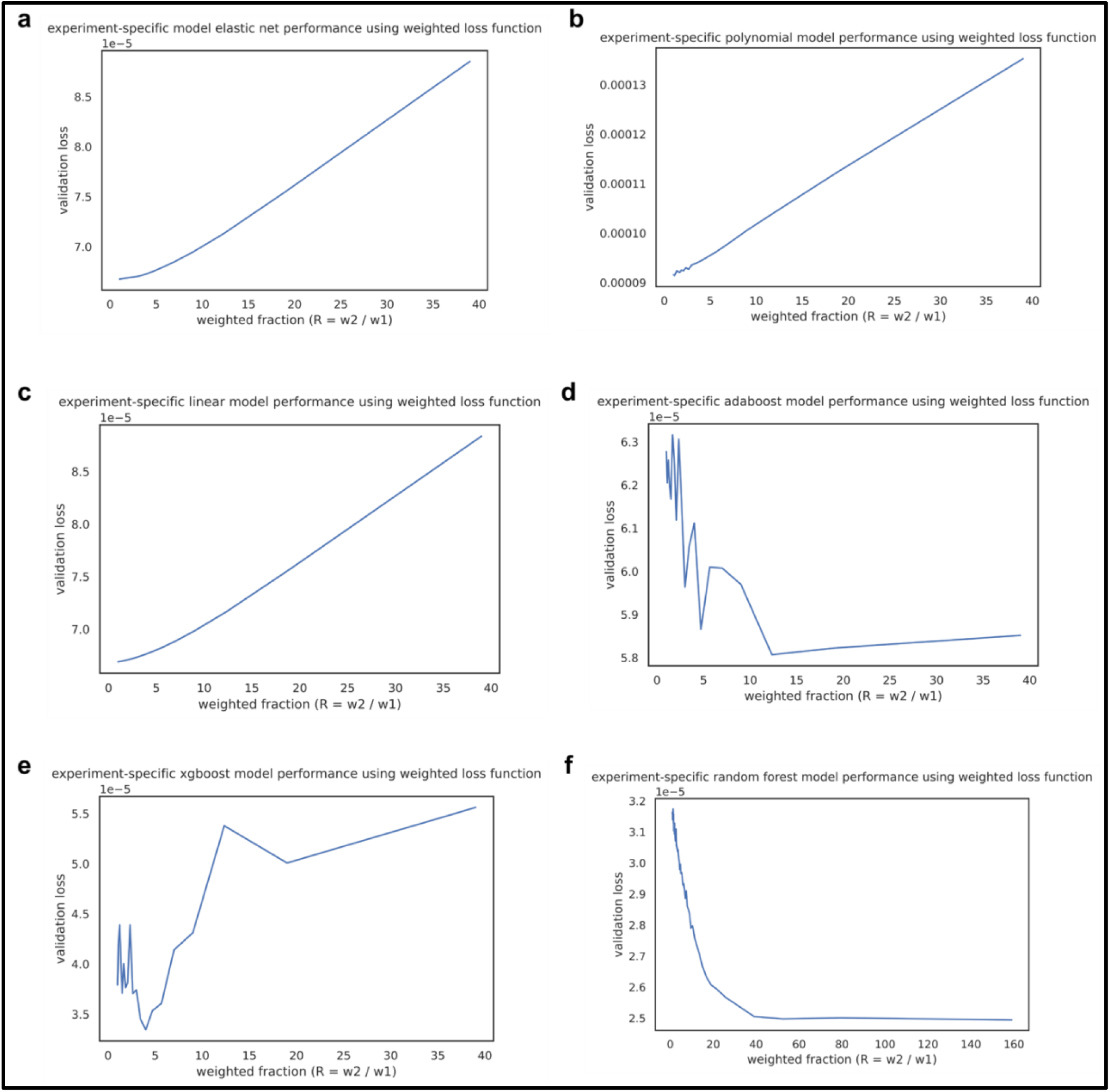
Hyperparameter tuning for weighted loss function. The hyperparameters, *w_1_* and *w_2_*, were tuned by setting both *w_1_* and *w_2_* to 0.5 initially and sequentially increasing the value of *w_2_* while decreasing the value of *w_1_* at each iteration, such that *w_1_* and *w_2_* always summed up to 1. This was performed under the hypothesis that training samples from the same experiment more accurately reflect the true dynamics of the experiment than those from other experiments. The example tuning curves were obtained from the optimally performing hybrid models in the rescue experiment under the discriminative approach.

**Figure S7:**
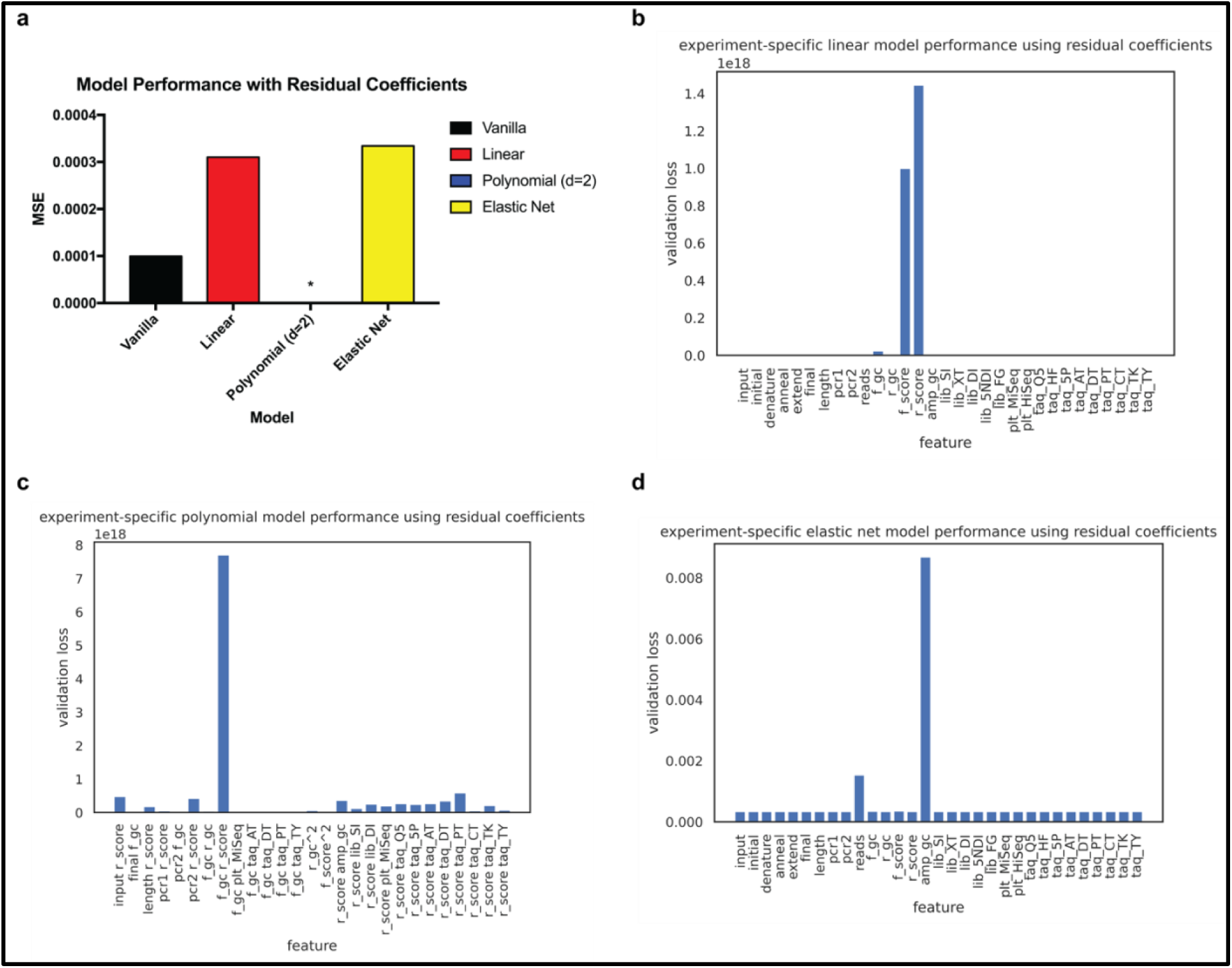
Model performances and optimizations for residual coefficient method. **a** LOOCV performances of discriminative hybrid models from the rescue experiment using the residual coefficient method. Polynomial model was omitted from the figure as they drastically overfitted on the training data as compared to the other models. **b-d** Validation losses of regression models measured by MSE when the corresponding coefficient *β_ι_* for each *feature_i_* in the linear regression function was sequentially tuned on experiment-specific SP samples.

**Table S1:**
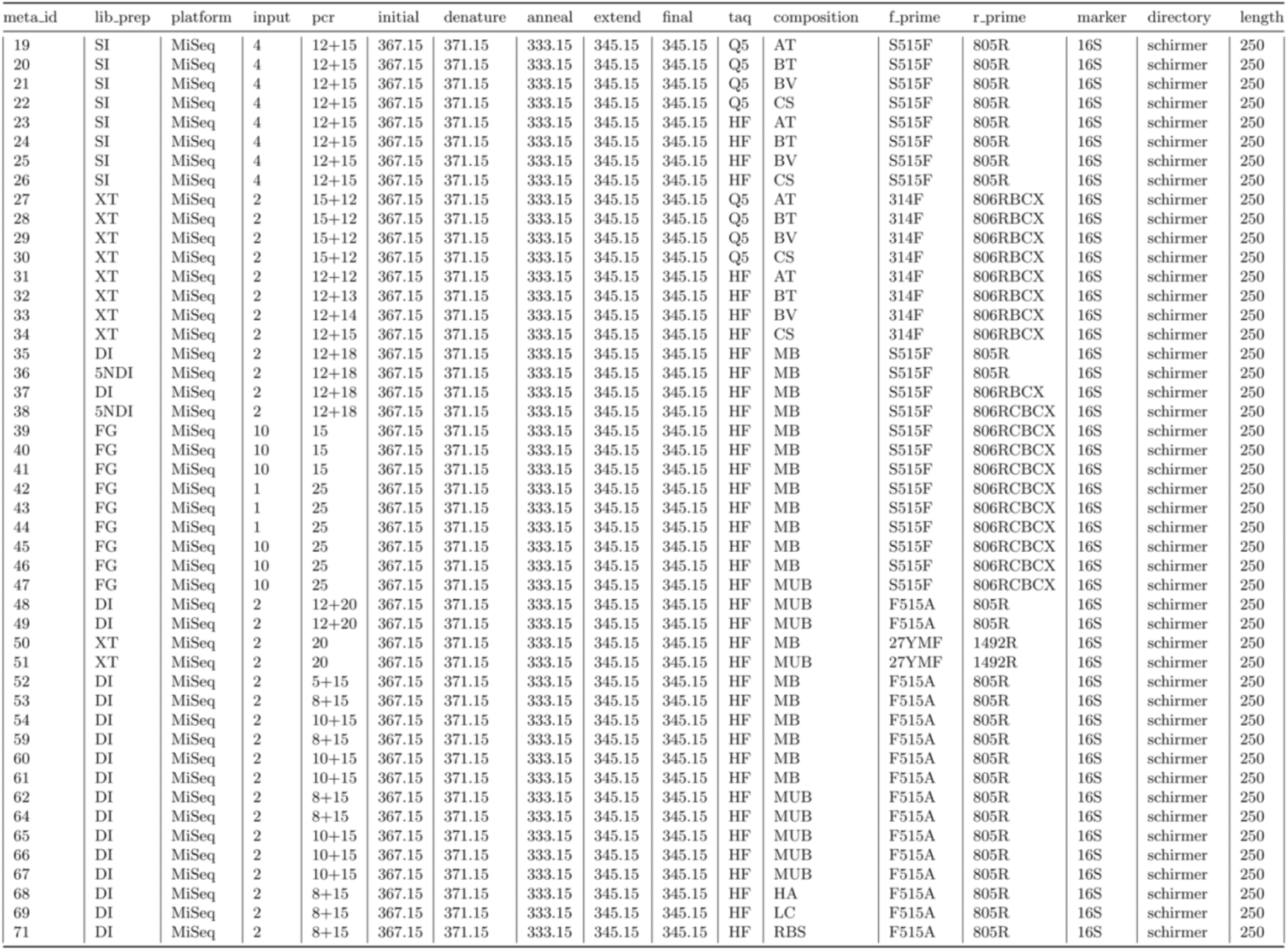
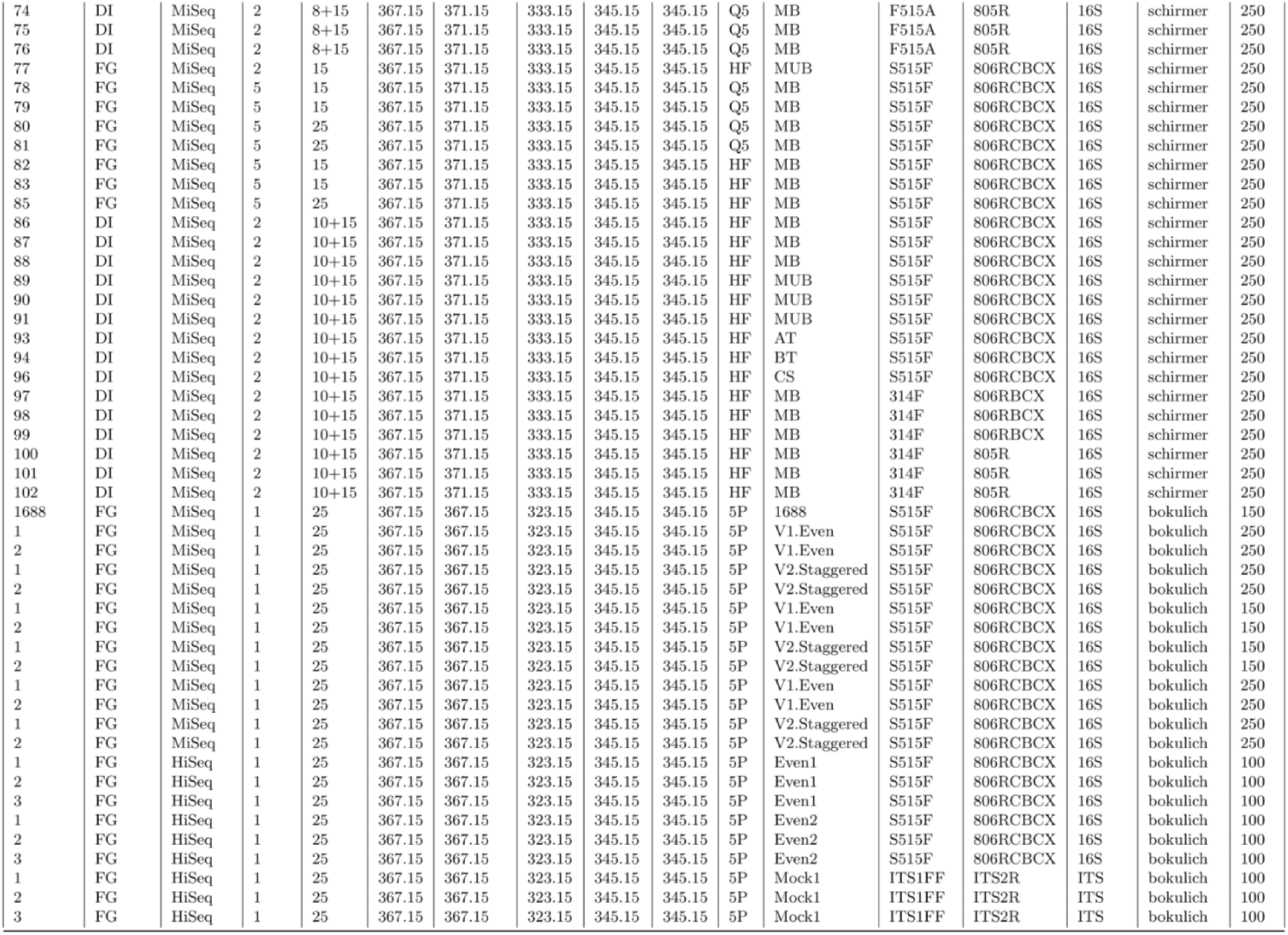
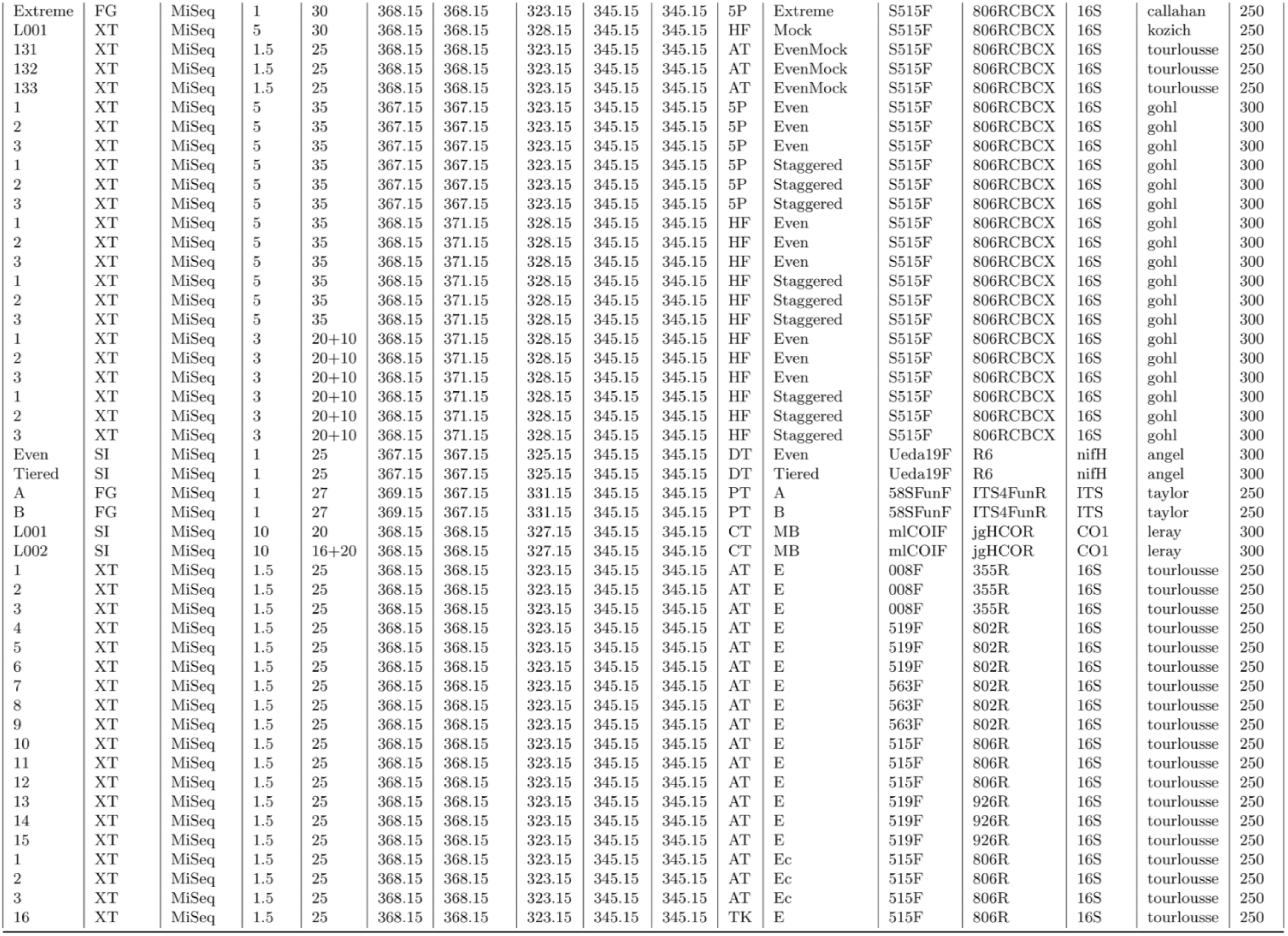
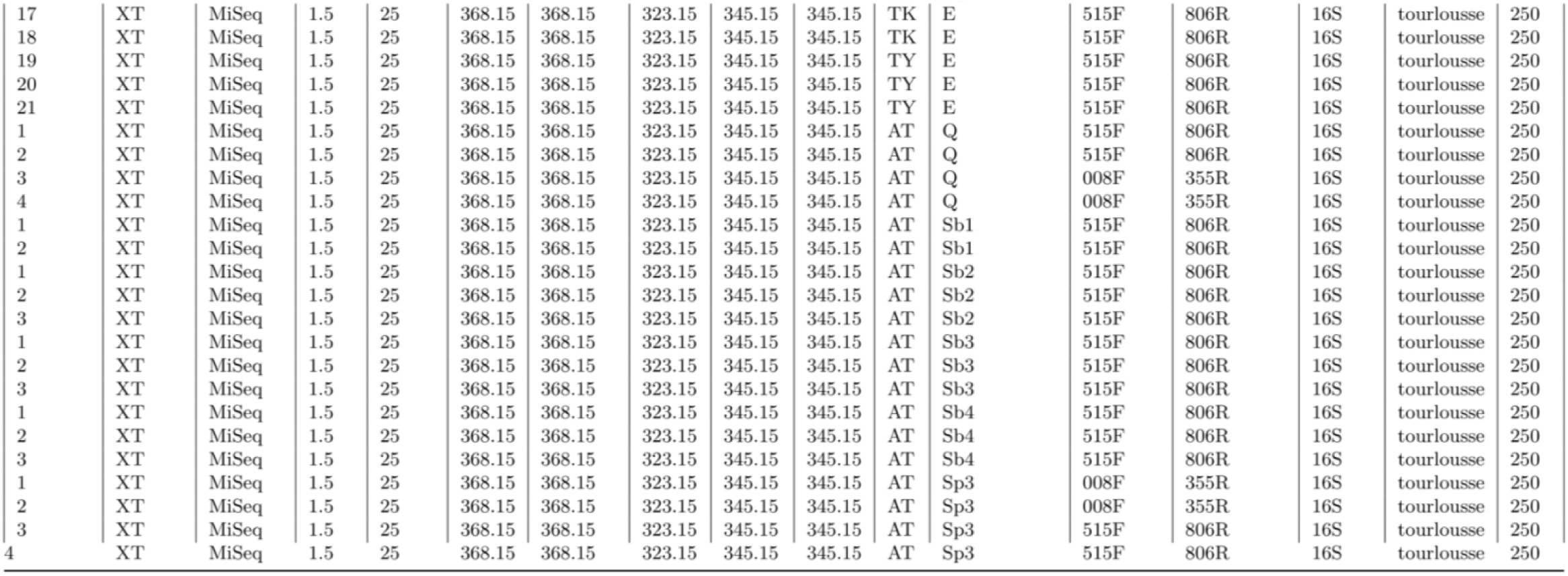
Overview of the experimental conditions extracted from the mock community datasets used in this study.

**Table S2:**
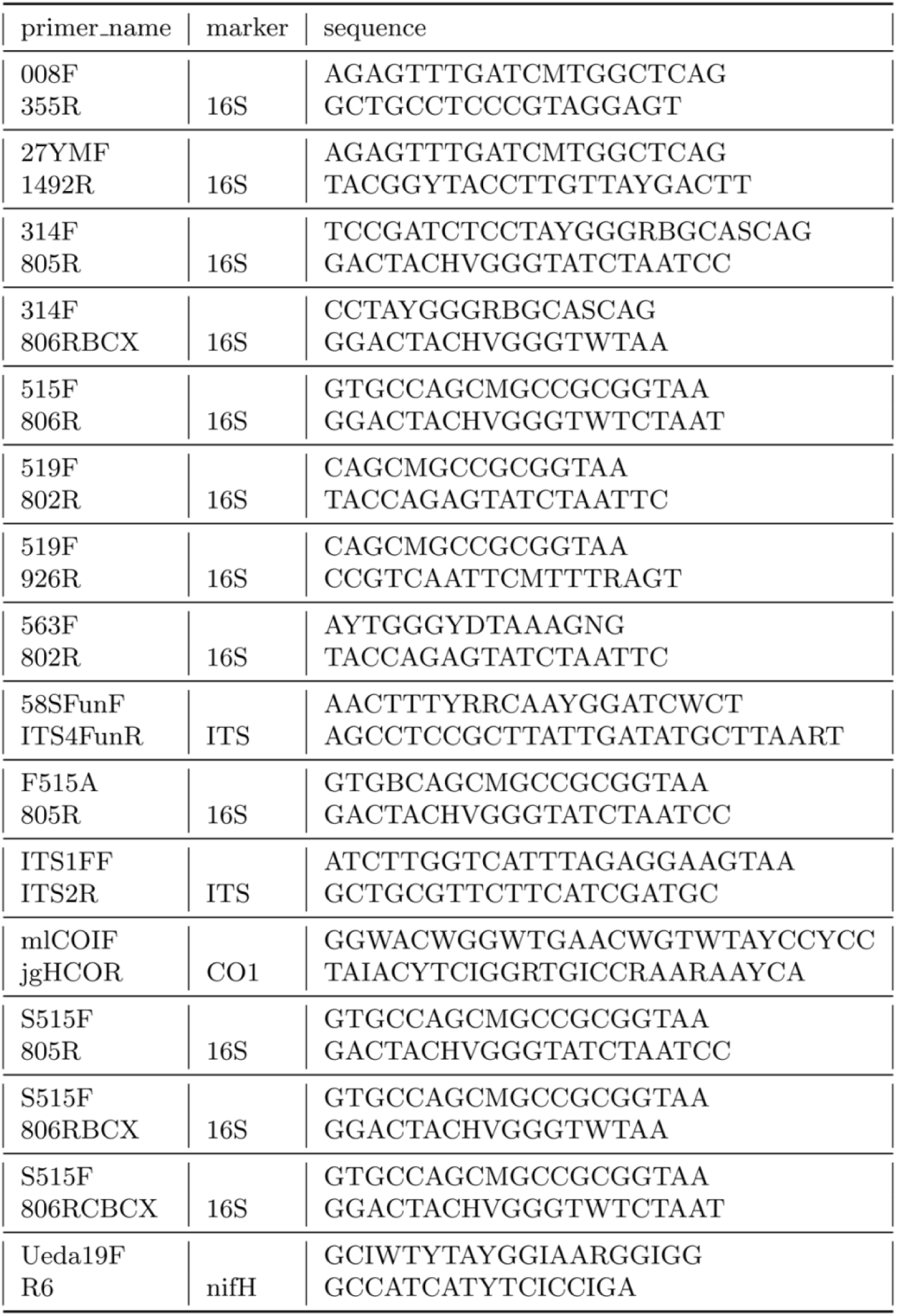
Sequences and markers of the PCR primer pairs used in this study.

**Table S3:** Identifiers of pseudo-taxa for synthetic spike-in constructs. Taxonomic identifiers that extend beyond the current maximum in the NCBI taxonomy database were assigned to the spike-in constructs prior to incorporating them as pseudo-taxa in the downstream metabarcoding workflow. The identifiers in the *spike_in* column were provided by Tourlousse et al. in the prior study (2017).

| taxid | spike_in |
| --- | --- |
| 2588930 | Ec5001 |
| 2588931 | Ec5002 |
| 2588932 | Ec5003 |
| 2588933 | Ec5004 |
| 2588934 | Ec5005 |
| 2588935 | Ec5501 |
| 2588936 | Ec5502 |
| 2588937 | Ec6001 |
| 2588938 | Bv5501 |
| 2588939 | Ca5501 |
| 2588940 | Ga5501 |
| 2588941 | Tb5501 |

## Notes

### Competing Interest Statement

The authors have declared no competing interest.

## References

Alan Agresti. Analysis of Ordinal Categorical Data. 2nd edition. Wiley; 2010. https://www.wiley.com/en-us/Analysis+of+Ordinal+Categorical+Data%2C+2nd+Edition-p-9780470082898. Accessed 27 Apr 2021.

Angel R, Nepel M, Panhölzl C, Schmidt H, Herbold CW, Eichorst SA, et al. Evaluation of Primers Targeting the Diazotroph Functional Gene and Development of NifMAP – A Bioinformatics Pipeline for Analyzing nifH Amplicon Data. Front Microbiol. 2018;9. doi:10.3389/fmicb.2018.00703.

Barnes MA, Turner CR. The ecology of environmental DNA and implications for conservation genetics. Conserv Genet. 2016;17:1–17.

Beentjes KK, Speksnijder AGCL, Schilthuizen M, Hoogeveen M, van der Hoorn BB. The effects of spatial and temporal replicate sampling on eDNA metabarcoding. PeerJ. 2019;7. doi:10.7717/peerj.7335.

Bokulich NA, Rideout JR, Kopylova E, Bolyen E, Patnode J, Ellett Z, et al. A standardized, extensible framework for optimizing classification improves marker-gene taxonomic assignments. PeerJ Inc.; 2015. doi:10.7287/peerj.preprints.934v2.

Brooks JP, Edwards DJ, Harwich MD, Rivera MC, Fettweis JM, Serrano MG, et al. The truth about metagenomics: quantifying and counteracting bias in 16S rRNA studies. BMC Microbiology. 2015;15:66.

Bru D, Martin-Laurent F, Philippot L. Quantification of the Detrimental Effect of a Single Primer-Template Mismatch by Real-Time PCR Using the 16S rRNA Gene as an Example. Appl Environ Microbiol. 2008;74:1660–3.

Buxton AS, Groombridge JJ, Zakaria NB, Griffiths RA. Seasonal variation in environmental DNA in relation to population size and environmental factors. Scientific Reports. 2017;7:46294.

Callahan BJ, McMurdie PJ, Rosen MJ, Han AW, Johnson AJA, Holmes SP. DADA2: High-resolution sample inference from Illumina amplicon data. Nature Methods. 2016;13:581–3.

Chen K, Hu Z, Xia Z, Zhao D, Li W, Tyler JK. The Overlooked Fact: Fundamental Need for Spike-In Control for Virtually All Genome-Wide Analyses. Molecular and Cellular Biology. 2016;36:662–7.

Chen T, Guestrin C. XGBoost: A Scalable Tree Boosting System. In: Proceedings of the 22nd ACM SIGKDD International Conference on Knowledge Discovery and Data Mining. New York, NY, USA: Association for Computing Machinery; 2016. p. 785–94. doi:10.1145/2939672.2939785.

Choo JM, Leong LE, Rogers GB. Sample storage conditions significantly influence faecal microbiome profiles. Scientific Reports. 2015;5:16350.

13. Claesen M, De Moor B. Hyperparameter Search in Machine Learning. arXiv:150202127 [cs, stat]. 2015. http://arxiv.org/abs/1502.02127. Accessed 28 Apr 2021.

D’Amore R, Ijaz UZ, Schirmer M, Kenny JG, Gregory R, Darby AC, et al. A comprehensive benchmarking study of protocols and sequencing platforms for 16S rRNA community profiling. BMC Genomics. 2016;17:55.

Deagle BE, Thomas AC, Shaffer AK, Trites AW, Jarman SN. Quantifying sequence proportions in a DNA-based diet study using Ion Torrent amplicon sequencing: which counts count? Molecular Ecology Resources. 2013;13:620–33.

Dietterich T, Kong EB. Machine Learning Bias, Statistical Bias, and Statistical Variance of Decision Tree Algorithms. Morgan Kaufmann Publishers Inc.; 1995. p. 313–21.

Djurhuus A, Closek CJ, Kelly RP, Pitz KJ, Michisaki RP, Starks HA, et al. Environmental DNA reveals seasonal shifts and potential interactions in a marine community. Nature Communications. 2020;11:254.

Dunn N, Priestley V, Herraiz A, Arnold R, Savolainen V. Behavior and season affect crayfish detection and density inference using environmental DNA. Ecol Evol. 2017;7:7777–85.

Evans NT, Olds BP, Renshaw MA, Turner CR, Li Y, Jerde CL, et al. Quantification of mesocosm fish and amphibian species diversity via environmental DNA metabarcoding. Molecular Ecology Resources. 2016;16:29–41.

Ficetola GF, Coissac E, Zundel S, Riaz T, Shehzad W, Bessière J, et al. An In silico approach for the evaluation of DNA barcodes. BMC Genomics. 2010;11:434.

Fisher A, Rudin C, Dominici F. Model Class Reliance: Variable Importance Measures for any Machine Learning Model Class, from the “Rashomon” Perspective. 2018. https://arxiv.org/abs/1801.01489v1. Accessed 28 Apr 2021.

Freund Y, Schapire RE. A Decision-Theoretic Generalization of On-Line Learning and an Application to Boosting. Journal of Computer and System Sciences. 1997;55:119–39.

Gohl DM, Vangay P, Garbe J, MacLean A, Hauge A, Becker A, et al. Systematic improvement of amplicon marker gene methods for increased accuracy in microbiome studies. Nature Biotechnology. 2016;34:942–9.

Hemery LG, Politano KK, Henkel SK. Assessing differences in macrofaunal assemblages as a factor of sieve mesh size, distance between samples, and time of sampling. Environ Monit Assess. 2017;189:413.

Highlander S. Mock Community Analysis. In: Nelson KE, editor. Encyclopedia of Metagenomics. New York, NY: Springer; 2013. p. 1–7. doi:10.1007/978-1-4614-6418-1_54-1.

Huber JA, Morrison HG, Huse SM, Neal PR, Sogin ML, Mark Welch DB. Effect of PCR amplicon size on assessments of clone library microbial diversity and community structure. Environ Microbiol. 2009;11:1292–302.

Jobson JD. Multiple Linear Regression. In: Jobson JD, editor. Applied Multivariate Data Analysis: Regression and Experimental Design. New York, NY: Springer; 1991. p. 219–398. doi:10.1007/978-1-4612-0955-3_4.

Jones MB, Highlander SK, Anderson EL, Li W, Dayrit M, Klitgord N, et al. Library preparation methodology can influence genomic and functional predictions in human microbiome research. PNAS. 2015;112:14024–9.

Kotsiantis SB. Supervised Machine Learning: A Review of Classification Techniques. In: Proceedings of the 2007 conference on Emerging Artificial Intelligence Applications in Computer Engineering: Real Word AI Systems with Applications in eHealth, HCI, Information Retrieval and Pervasive Technologies. NLD: IOS Press; 2007. p. 3–24.

Kozich JJ, Westcott SL, Baxter NT, Highlander SK, Schloss PD. Development of a Dual-Index Sequencing Strategy and Curation Pipeline for Analyzing Amplicon Sequence Data on the MiSeq Illumina Sequencing Platform. Appl Environ Microbiol. 2013;79:5112–20.

Krehenwinkel H, Fong M, Kennedy S, Huang EG, Noriyuki S, Cayetano L, et al. The effect of DNA degradation bias in passive sampling devices on metabarcoding studies of arthropod communities and their associated microbiota. PLOS ONE. 2018;13:e0189188.

Krehenwinkel H, Wolf M, Lim JY, Rominger AJ, Simison WB, Gillespie RG. Estimating and mitigating amplification bias in qualitative and quantitative arthropod metabarcoding. Scientific Reports. 2017;7:17668.

Kumar A, Kaur J. Primer Based Approach for PCR Amplification of High GC Content Gene: Mycobacterium Gene as a Model. Molecular Biology International. 2014;2014:e937308.

Lancichinetti A, Fortunato S, Kertesz J. Detecting the overlapping and hierarchical community structure of complex networks. New J Phys. 2009;11:033015.

Leray M, Knowlton N. Random sampling causes the low reproducibility of rare eukaryotic OTUs in Illumina COI metabarcoding. PeerJ. 2017;5:e3006.

Loos LM van der, Nijland R. Biases in bulk: DNA metabarcoding of marine communities and the methodology involved. Molecular Ecology. n/a n/a. 10.1111/mec.15592.

Ma X, Shao Y, Tian L, Flasch DA, Mulder HL, Edmonson MN, et al. Analysis of error profiles in deep next-generation sequencing data. Genome Biol. 2019;20:50.

Matesanz S, Pescador DS, Pías B, Sánchez AM, Chacón-Labella J, Illuminati A, et al. Estimating belowground plant abundance with DNA metabarcoding. Molecular Ecology Resources. 2019;19:1265–77.

Nichols RV, Vollmers C, Newsom LA, Wang Y, Heintzman PD, Leighton M, et al. Minimizing polymerase biases in metabarcoding. Molecular Ecology Resources. 2018;18:927–39.

Pawluczyk M, Weiss J, Links MG, Egaña Aranguren M, Wilkinson MD, Egea-Cortines M. Quantitative evaluation of bias in PCR amplification and next-generation sequencing derived from metabarcoding samples. Anal Bioanal Chem. 2015;407:1841–8.

Piñol J, Senar MA, Symondson WOC. The choice of universal primers and the characteristics of the species mixture determine when DNA metabarcoding can be quantitative. Molecular Ecology. 2019;28:407–19.

Pinto AJ, Raskin L. PCR Biases Distort Bacterial and Archaeal Community Structure in Pyrosequencing Datasets. PLOS ONE. 2012;7:e43093.

Polz MF, Cavanaugh CM. Bias in Template-to-Product Ratios in Multitemplate PCR. Appl Environ Microbiol. 1998;64:3724–30.

Schirmer M, Ijaz UZ, D’Amore R, Hall N, Sloan WT, Quince C. Insight into biases and sequencing errors for amplicon sequencing with the Illumina MiSeq platform. Nucleic Acids Res. 2015;43:e37.

Sigsgaard EE, Nielsen IB, Carl H, Krag MA, Knudsen SW, Xing Y, et al. Seawater environmental DNA reflects seasonality of a coastal fish community. Mar Biol. 2017;164:128.

Takahara T, Minamoto T, Yamanaka H, Doi H, Kawabata Z. Estimation of Fish Biomass Using Environmental DNA. PLOS ONE. 2012;7:e35868.

Taylor DL, Walters WA, Lennon NJ, Bochicchio J, Krohn A, Caporaso JG, et al. Accurate Estimation of Fungal Diversity and Abundance through Improved Lineage-Specific Primers Optimized for Illumina Amplicon Sequencing. Appl Environ Microbiol. 2016;82:7217–26.

Tourlousse DM, Yoshiike S, Ohashi A, Matsukura S, Noda N, Sekiguchi Y. Synthetic spike-in standards for high-throughput 16S rRNA gene amplicon sequencing. Nucleic Acids Res. 2017;45:e23.

Vasselon V, Bouchez A, Rimet F, Jacquet S, Trobajo R, Corniquel M, et al. Avoiding quantification bias in metabarcoding: Application of a cell biovolume correction factor in diatom molecular biomonitoring. Methods in Ecology and Evolution. 2018;9:1060–9.

Wagner Mackenzie B, Waite DW, Taylor MW. Evaluating variation in human gut microbiota profiles due to DNA extraction method and inter-subject differences. Front Microbiol. 2015;6. doi:10.3389/fmicb.2015.00130.

Wood DE, Lu J, Langmead B. Improved metagenomic analysis with Kraken 2. Genome Biology. 2019;20:257.

Zhang Z, Lai Z, Xu Y, Shao L, Wu J, Xie G-S. Discriminative Elastic-Net Regularized Linear Regression. IEEE Transactions on Image Processing. 2017;26:1466–81.

